# Loss of NAT10 disrupts enhancer organization via p300 mislocalization and suppresses transcription of genes necessary for metastasis progression

**DOI:** 10.1101/2024.01.24.577116

**Authors:** Ruhul Amin, Ngoc-Han Ha, Tinghu Qiu, Ronald Holewinski, Khiem C. Lam, Amélie Lopès, Huaitian Liu, Andy D. Tran, Maxwell P. Lee, Supuni Thalalla Gamage, Thorkell Andresson, Romina S. Goldszmid, Jordan L. Meier, Kent W. Hunter

## Abstract

Acetylation of protein and RNA represent a critical event for development and cancer progression. NAT10 is the only known RNA acetylase that catalyzes the N4-actylcytidine (ac4C) modification of RNAs. Here, we show that the loss of NAT10 significantly decreases lung metastasis in allograft and genetically engineered mouse models of breast cancer. NAT10 interacts with a mechanosensitive, metastasis susceptibility protein complex at the nuclear pore. In addition to its canonical role in RNA acetylation, we find that NAT10 interacts with p300 at gene enhancers. NAT10 loss is associated with p300 mislocalization into heterochromatin regions. NAT10 depletion disrupts enhancer organization, leading to alteration of gene transcription necessary for metastatic progression, including reduced myeloid cell-recruiting chemokines that results in a less metastasis-prone tumor microenvironment. Our study uncovers a distinct role of NAT10 in enhancer organization of metastatic tumor cells and suggests its involvement in the tumor-immune crosstalk dictating metastatic outcomes.

## Introduction

Metastasis, a complex and devastating process by which cancer cells disseminate from their primary site to distant organs, remains one of the most formidable challenges in the field of oncology^1, 2^. Metastasis is responsible for the majority of cancer-related deaths and significantly limits the success of existing therapeutic strategies^3^. The intricate interplay of numerous genetic^1, 4^, epigenetic^5^, and microenvironmental factors^6^ has been implicated in the metastatic cascade, underscoring the need for a comprehensive understanding of the underlying molecular mechanisms.

Our previous research has shown that individual genetic polymorphisms play a substantial role in determining the metastatic potential of tumor cells^7^. Since the majority of these inherited variations are found in noncoding regions of the DNA, they are believed to impact metastasis by modifying gene expression rather than directly affecting protein function. By performing genome-wide analysis of metastasis-associated polymorphisms in noncoding regulatory regions, we have shown that polymorphisms in the promoter region of genes can predict metastasis outcomes in both human estrogen receptor (ER) positive^8^ and ER negative^9^ breast cancer patients. Our analysis has led to the identification of several metastasis susceptibility genes, including *Sipa1*^10^, *Rrp1b*^11^, the short isoform of *Brd4* (Brd4-SF)^12^, *Arntl2*^9^ and *Nup210*^8^. Interestingly, many of these factors exhibit a commonality in their interaction at the nuclear lamina or nuclear pore^8, 13^. NUP210, a nuclear pore associated protein has been shown to facilitate cellular mechanosensation in metastatic tumor cells. Since mechanical signals from the extracellular microenvironment appear to be critical for tumor and metastatic progression^14^, we sought to investigate the role of these proteins in cellular mechanosensation and metastasis. Our previous mass spectrometry analysis revealed that BRD4-SF also interacts with NAT10 at the nuclear lamina^13^. In addition, *in vitro* promoter pull down followed by mass spectrometry analysis revealed that NAT10 differentially bind to the promoters of high-versus low-metastasis Arntl2 alleles^9^. However, individual function of NAT10 in cellular mechanosensation and metastasis was not defined. We therefore selected NAT10 for further study to investigate its role in metastasis susceptibility.

NAT10, a ubiquitously expressed enzyme, functions as an RNA cytidine acetyl transferase (ac4C) for mRNAs^15^, rRNAs and tRNAs^16^. Recently, it has gained recognition as a key modulator of various cellular processes. Some reports suggest its involvement in protein acetylation^17–19^. The role of NAT10 in cancer has garnered increasing interest, with mounting evidence implicating it in both cancer development^20–23^ and metastatic progression^24–27^. However, most studies have focused on acetylation of individual substrates by NAT10. In this study, we employed an unbiased approach that combines genomics, proteomics, and transcriptomics to explore the role of NAT10 in metastasis. Our findings reveal a previously unknown mechanism by which NAT10 contributes to metastasis through p300-associated chromatin remodeling, which appears to be separate from its traditional function in RNA acetylation, ultimately enhancing the metastatic potential of tumor cells and their crosstalk with innate immune myeloid cells to form a metastasis-promoting primary tumor microenvironment.

## Materials and methods

### Mouse strains

The animals utilized in this investigation were subjected to the guidelines outlined in animal study protocols LPG-002 and LCBG-004, approved by the National Cancer Institute (NCI) at the Bethesda Animal Use and Care Committee. Euthanasia of the animals was carried out through anesthesia using Avertin injection, followed by cervical dislocation. Female mice of the BALB/c (Strain #: 000651) and FVB/NJ (Strain #: 001800) strains were procured from The Jackson Laboratory. Nat10 mutant mice C57BL/6-Nat10^H537A^ were generated by Dr. Jordan L. Meier’s lab at the NCI. MMTV-PyMT (FVB/N-Tg(MMTV-PyVT)634Mul/J, Strain #: 002374) mice were purchased from The Jackson Laboratory.

### Cell culture

Dr. Lalage Wakefield (NCI, NIH) provided the 4T1 and 6DT1 mouse mammary tumor cell lines. These cell lines were cultured in Dulbecco’s modified Eagle’s medium (DMEM) (Gibco), supplemented with 9% fetal bovine serum (FBS) (Gemini), 1% L-glutamine (Gibco), and 1% penicillin–streptomycin (Gemini). Human 293FT cells, acquired from Thermo Fisher Scientific, were cultured using the same medium mentioned above.

### Cloning

cDNAs encoding mouse and human NAT10 were purchased from Horizon Discovery. Human and mouse *Nat10* coding regions were first cloned into Gateway entry vector pENTR-DTOPO. These entry vectors were then subsequently combined with entry vectors containing the pol2 promoter and Flag/Myc-tag into a custom lentiviral destination vector pDest-659 (a gift from Dominic Esposito, NCI) using Gateway LR Clonase (Thermo Fisher Scientific) to generate the final *Nat10* overexpression vector. 3X Flag human *BRD4* short or long isoform vectors were generated in the Protein Expression Laboratory of Dr. Dominic Esposito at the Frederick National Laboratory for Cancer Research, NCI.

### Lentiviral production and generation of stable cell lines

TRC lentiviral shRNA vectors were purchased from Horizon Discovery. For mouse Nat10 shRNA knockdown, sh-Nat10-E8 (TRCN0000184285: ATACAGATGGAACGATGGTGC), sh-Nat10-E10 (TRCN0000184551: ATTAAGCCACTTCTCCACTGC), and sh-Nat10-E12 (TRCN0000179570: TTCAAGAGTGACTTTGTCTTC) were used. To produce lentivirus, plasmids containing shRNA, along with packaging plasmids psPAX2 (Addgene plasmid #12260) and envelope plasmid pMD2.G (Addgene plasmid #12259), generously provided by the Trono lab, were transfected into the human 293FT cell line using X-tremeGENE 9 DNA transfection reagent (Roche). After 48 hours, the culture supernatant, which contained lentivirus, was collected, filtered through a 0.45 μm filter (Millipore), and then utilized for transducing mouse mammary tumor cell lines. shRNAs that stably integrated into mouse cells were chosen through selection with 10 μg/ml puromycin (Sigma). For the generation of Nat10-overexpressing cells, similar lentiviral packaging was performed using lentiviral plasmids carrying mouse *Nat10* coding sequence. Following transduction, Nat10-expressing cells were selected using Blasticidin (Gibco) at a concentration of 10 μg/ml.

### CRISPR/Cas9-mediated knockout of the mouse *Nat10* gene

To perform knockout of *Nat10* in mouse 4T1 cells, the Cas9 D10A double-nicking strategy was employed. Sense (5ʹ-TGGTGTTATAAGAAAGAGCT-3ʹ) and antisense (5ʹ-CCGAGCCTTCACAGTTGCCT-3ʹ) sgRNAs spanning exon 3 of *Nat10* were designed using the CRISPic web portal of the Broad Institute (https://portals.broadinstitute.org/gppx/crispick/public). The All-in-One plasmid, which encodes dual U6 promoter-driven sgRNAs and an EGFP-coupled Cas9 D10A plasmid (AIO-GFP), was provided by Dr. Steve Jackson (The Wellcome Trust Sanger Institute, UK) (Addgene #74119). The sgRNAs were inserted into the AIO-GFP plasmid using the method described previously^28^. In brief, the AIO-GFP plasmid was first digested with the BbsI restriction enzyme, dephosphorylated using calf intestinal phosphatase, and gel purified. sgRNA oligonucleotide pairs were obtained from Integrated DNA Technologies, annealed, and phosphorylated with T4 polynucleotide kinase (NEB). First, the sgRNA was cloned into the BbsI site, and second, the sgRNA was cloned into the BsaI site. The final clone’s DNA sequence was confirmed through Sanger sequencing using a specific primer (5′-CTTGATGTACTGCCAAGTGGGC-3′). The sgRNA plasmid was then transfected into 4T1 cells using Nanojuice transfection reagent (Millipore). After 48 hours, GFP-positive cells were sorted using flow cytometry, and single cells were plated on a 96-well plate. Cas9 D10A-edited clones were identified through PCR and Sanger sequencing of genomic DNA.

### Generation of NAT10-dTAG knockin cell line using the CRISPR/Cas9 method

The NAT10-dTAG knockin vector was constructed using the protocol described elsewhere^29^. C-terminal targeting guide RNA was designed using CRISPick webpage. sgRNA oligonucleotides purchased from IDT were annealed and ligated in the BbsI restriction site of pX330A-1×2 (Addgene, #58766) plasmid to make pX330A-1x2-cNAT10 vector. PITCh sgRNA from pX330S-2-PITCh (Addgene #63670) vector was then inserted into the pX330A-1x2-cNAT10 vector using Golden Gate Assembly (New England Biolab) to make the pX330A-1x2-cNAT10/PITCh vector. For making donor vector containing dTAG-2xHA sequences, the pCRIS-PITChv2-dTAG-Puro (BRD4) (Addgene, #91796) vector was used. This plasmid was used as a template for PCR amplification of the dTAG-2xHA sequence. PCR was performed using primers containing the homology arm flanking the sgRNA region (replacing the BRD4 homology arm) and Gibson cloned (NEBuilder Hifi DNA assembly kit, NEB) into the same vector by digesting with MluI restriction enzyme.

For the generation of the NAT10 dTAG knockin cell line, human 293FT cells were grown on 10cm dishes and transfected with sgRNA-containing vector (pX330A-1x2-cNAT10/PITCh) and donor vector using Xtremegene 9 transfection reagent (Roche). 72 h after transfection, positive clones containing the c-terminal NAT10 dTAG knockin were selected with 1 μg/ml puromycin for 7 days and subjected to single-cell cloning. NAT10 knockin was confirmed by PCR genotyping. Further validation of knockin clones was performed using western blotting with an anti-HA tag antibody. For the induction of NAT10 degradation, 1 μM dTAG-13 (Tocris Bioscience) was used for 24 h.

### Spontaneous metastasis assays

Female BALB/c and FVB/NJ mice, aged around 6 weeks, were acquired from The Jackson Laboratory. To perform orthotopic transplantation of cells with gene knockdown or overexpression, 100,000 cells were injected into the fourth mammary fat pad of the mice. After 30–35 days, the mice were euthanized, primary tumors were removed, weighed, and the surface lung metastases were quantified. For the autochthonous genetically engineered model, female C57BL/6-Nat10^H537A^ mice were crossed with male MMTV-PyMT mice. After the pups were born, genotyping was performed using tail biopsy DNA. Wild type and heterozygous *Nat10* mutant mice with PyMT antigen positive pups were screened and allowed to develop tumors. After ∼120 days, mice were euthanized, all the primary tumors were collected, weighted, and the surface lung metastases were counted.

### Protein lysate preparation and western blotting

Whole-cell protein lysate was prepared by utilizing lysis buffer containing 20 mM Tris-HCl pH 8.0, 400 mM NaCl, 5 mM EDTA, 1 mM EGTA, 10 mM NaF, 1 mM sodium pyrophosphate, 1% Triton X-100, 10% glycerol, and a protease and phosphatase inhibitor cocktail. For the preparation of nuclear protein lysates, the Nuclear Complex Co-IP Kit (Active Motif) was used according to manufacturer’s instruction. The protein concentration was determined using the Pierce BCA Protein Assay Kit (Thermo Fisher Scientific). Subsequently, 20-40 μg of protein lysates were combined with 4X NuPAGE LDS sample buffer (Invitrogen) and 10X NuPAGE Sample Reducing Agent (Invitrogen), followed by boiling at 95 °C for 5 min. The resolved samples were run on either NuPAGE 3–8% Tris-acetate or NuPAGE 12% Bis-Tris protein gels (Thermo Fisher Scientific) with the appropriate running buffer. Proteins were then transferred onto a PVDF membrane (Millipore), and the membrane was blocked with blocking buffer (TBST + 5% non-fat dry milk) for 1 h. Subsequent steps involved overnight incubation with the relevant primary antibodies, washing with TBST, incubation with secondary antibodies for 1 h, and development of the signal on X-ray film using the Amersham ECL Prime Western Blotting Detection Reagent (Cytiva).

Following antibodies and dilutions were used: rabbit NAT10 (1:5000, Abcam), mouse Nat10 (1:5000, Proteintech), mouse β-actin (1:10,000; Abcam), rabbit NUP210 (1:1000, Fortis Life Sciences), rabbit SIPA1 (1:1000, Abcam), rabbit pan-BRD4 (1:5000, Cell Signaling Technology), mouse Flag-tag (1:5000, Millipore-Sigma), rabbit Flag-tag (1:1000, Cell Signaling Technology), Mouse Myc-tag (1:5000, Cell Signaling Technology), rabbit ITGB1 (1:1000, Cell Signaling Technology), mouse H3.1/3.2 (1:5000, Active Motif), rabbit RRP1B (1:5000, Millipore-Sigma), mouse NPM1 (1:1000, Abcam), rabbit α/β-tubulin (1:5000, Cell Signaling Technology), rabbit H3.3 (1:5000, Abcam), mouse pan-H4 (1:1000, Cell Signaling Technology), rabbit H4K12ac (1:5000, Cell Signaling Technology), rabbit H4K20ac (1:5000, Millipore-Sigma), rabbit H4K5ac (1:1000, Active Motif), rabbit H4K8ac (1:1000, Abcam), rabbit H3K27ac (1:1000, Cell Signaling Technology), mouse pan-H3 (1:10000, Cell Signaling Technology), rabbit H4K16ac (1:1000, Cell Signaling Technology), rabbit pan-H4ac (1:1000, Active Motif), rabbit pan-ac-K (1:1000, Cell Signaling Technology), Rabbit p300 (1:5000, Cell Signaling Technology), rabbit ac-p300 (K1499)/ac-CBP (K1535) (1:5000, Cell Signaling Technology), rabbit phospho-p300 (S89) (1:300, Aviva), rabbit c-MYC (1:5000, Cell Signaling Technology), rabbit EIF2α (1:1000, Cell Signaling Technology), rabbit phospho-EIF2α (1:1000, Cell Signaling Technology), rabbit p70 S6 Kinase (1:1000, Cell Signaling Technology), rabbit phospho-p70 S6 Kinase (T421/S424) (1:1000, Cell Signaling Technology). HRP-conjugated mouse secondary antibody (GE Healthcare) was used at 1:10,000 dilution. HRP-conjugated rabbit secondary antibody (Cell Signaling Technology) was used at 1:5,000 dilution.

### Co-immunoprecipitation

The Nuclear Complex Co-IP Kit (Active Motif) was utilized for co-immunoprecipitation on nuclear lysates. 4T1 cells were seeded onto 15 cm tissue culture dishes with a seeding density of 4 × 10^6^ cells per dish. After 48 h of incubation, cells were harvested, and nuclear lysates were prepared. Subsequently, 200–500 μg of nuclear lysates were incubated with 2 μg of specific antibodies and 50 μg of Dynabeads Protein G (Invitrogen). Following an overnight incubation on a rotator at 4 °C, immune complexes were isolated using a magnetic stand. The beads were then washed 3 times, resuspended in 2X NuPAGE LDS sample buffer (Invitrogen), and incubated at 95 °C in a heat block for 5 min. Finally, the samples were loaded onto NuPAGE protein gels, and the standard western blot protocol was followed as described previously.

### Transfection

For the coimmunoprecipitation, human 293FT cells were grown on 15 cm tissue culture dishes at a seeding density of 5 x 10^6^ cells per dish. 24 h later, cells were co-transfected with 5 μg of Myc-tagged NAT10 and 5 μg of BRD4 short or long isoform using the Xtremegene 9 DNA transfection reagent (Roche). After overnight incubation, transfection medium was replaced by fresh medium. After 24 h of incubation, cells were harvested, and nuclear lysates were prepared using the Nuclear Complex Co-IP kit (Active Motif).

### Digestion, TMTpro labeling, and LC/MS analysis

#### Digestion of Nat10 IP proteins and TMTpro labeling

4T1 nuclear lysates were prepared using the Nuclear Complex Co-IP kit (Active Motif) according to the manufacturer’s instruction. Each sample bound to capture beads was resuspended in 200 μl of EasyPep lysis buffer, 50 μl reducing buffer, and 50 μl alkylating solution provided with the EasyPep kit (Thermo Fisher Scientific). Samples were incubated at 25°C for 1 h in the dark at 1000 rpm then 50 μl of 100 ng trypsin/LysC was added. Samples were further incubated overnight for 23 h total at 37°C with shaking at 1000rpm. After digestion 200 μl of each solution was transferred to a new tube and treated with 50 µl of 2 μg/μl TMTpro (Thermo Fisher Scientific) reagent, then incubated for 1 h at 25⁰C with shaking. Excess TMTpro was quenched with 50 μl of 5% hydroxylamine, 20% formic acid for 10 min, then samples were combined. Samples were cleaned using EasyPep mini columns (Thermo Fisher Scientific) as described in the manual. Eluted peptides were dried in a speed-vac.

#### Digestion of 4T1 nuclear lysates, TMTpro labeling, and acetylation enrichment

4T1 nuclear lysates were prepared as described. Proteins concentration was determined by BCA and 120 µg of each sample was precipitated by the addition of TCA (1:4 v/v ratio of TCA/sample).Next, 200 µl of EasyPep lysis buffer containing 1X deacetylase inhibitors (Active Motif) and 40 ng/µl trypsin/LysC and 50 µl each of reducing and alkylation solutions (provided with the EaspPep kit) were added to each protein pellet. Samples were then incubated at 37°C for 18 h with shaking at 800 rpm. TMTpro labels were added (750µg) to each sample and incubated for 1 h at 25 ⁰C with shaking. Excess TMTpro was quenched with 50 μl of 5% hydroxylamine, 20% formic acid for 10 min and samples were then combined and cleaned using EasyPep Maxi (Thermo Fisher Scientific) column as described in the manual. Peptides were eluted in 3 ml and 25 µl was dried and saved for total protein analysis. The rest of the eluate was dried and enriched for acetylated peptides using the PTMScan® HS Acetyl-Lysine Motif (Ac-K) Kit (Cell Signaling Technology) as described in the manual. Peptides were eluted 2x with 200 µl of 50%ACN, 0.5%TFA and eluted fractions were combined and dried.

#### LC/MS analysis of Nat10-IP peptides

Peptides were resuspended in 50 μl of 0.1% FA and 5 μl was analyzed in triplicate using a Dionex U3000 RSLC in front of an Orbitrap Eclipse (Thermo Fisher Scientific) equipped with an EasySpray ion source. Solvent A consisted of 0.1%FA in water and Solvent B consisted of 0.1%FA in 80%ACN. The loading pump contained Solvent A and was operated at 7 μl per min for the first 6 minutes of the run then dropped to 2 μl per min when the valve was switched to bring the trap column (Acclaim™ PepMap™ 100 C18 HPLC Column, 3 μm, 75 μm I.D., 2 cm, PN 164535) in-line with the analytical column EasySpray C18 HPLC Column, 2 μm, 75 μm I.D., 25 cm, PN ES902). The gradient pump was operated at a flow rate of 300 nl per min. Each run used a linear LC gradient of 5-7%B for 1min, 7-30%B for 83min, 30-50%B for 25 min, 50-95%B for 4 min, holding at 95%B for 7 min, then re-equilibration of analytical column at 5%B for 17 min. All MS injections employed the TopSpeed method with three FAIMS compensation voltages (CVs) and a 1 s cycle time for each CV (3 s cycle time total) that consisted of the following: spray voltage of 2200V and ion transfer temperature of 300⁰C. MS1 scans were acquired in the Orbitrap with a resolution of 120,000, AGC of 4 x 10^5^ ions, and max injection time of 50ms, mass range of 350-1600 m/z; MS2 scans were acquired in the Orbitrap using the TurboTMT method with a resolution of 15,000, AGC of 1.25 x 10^5^, max injection time of 22 ms, HCD energy of 38%, isolation width of 0.4Da, intensity threshold of 2.5 x 10^4^ and charges 2-6 for MS2 selection. Advanced Peak Determination, Monoisotopic Precursor selection (MIPS), and EASY-IC for internal calibration were enabled and dynamic exclusion was set to a count of 1 for 15 s. The only difference in the methods was the CVs used; one method used CVs of -45, -60, -75, the second used CVs of -50, -65, -80, and the third used CVs of -55, -70, -85.

#### LC/MS analysis of 4T1 nuclear global and acetyl peptides

Enriched acetylated peptides were resuspended in 20 μl of 0.1% FA and 15 μl was loaded once, while a global protein aliquot was resuspended in 50 μl of 0.1% FA and 5 µl was loaded twice onto a Dionex U3000 RSLC in front of an Orbitrap Eclipse (Thermo Fisher) equipped with an EasySpray ion source. Solvent A consisted of 0.1%FA in water and Solvent B consisted of 0.1%FA in 80%ACN. The loading pump contained Solvent A and was operated at 7 μL/min for the first 6 minutes of the run then dropped to 2 μl/min when the valve was switched to bring the trap column (Acclaim™ PepMap™ 100 C18 HPLC Column, 3 μm, 75 μm I.D., 2 cm, PN 164535) in-line with the analytical column EasySpray C18 HPLC Column, 2 μm, 75 μm I.D., 25 cm, PN ES902). The gradient pump was operated at a flow rate of 300 nl/min. Acetylated peptides used a linear LC gradient of 5-7%B for 1 min, 7-30%B for 33 min, 30-50%B for 15 min, 50-95%B for 4 min, holding at 95%B for 7 min, then re-equilibration of analytical column at 5%B for 17 min. Total protein analysis used a linear LC gradient of 5-7%B for 1 min, 7-30%B for 133 min, 30-50%B for 35 min, 50-95%B for 4 min, holding at 95%B for 7 min, then re-equilibration of the analytical column at 5%B for 17 min. All MS injections employed the TopSpeed method with a 3 s cycle time. Acetylated peptide analysis used three FAIMS compensation voltages (CVs) of -45, -60, -75 and a 1 s cycle time for each CV. Total peptide analysis used four CVs of -45, -55, -65, -75 for one method and -50, -60, -70, -80 for the second method and a 0.75 s cycle time for each CV. For all methods the spray voltage was 2200 V and the ion transfer temperature was 300⁰C. MS1 scans were acquired in the Orbitrap with resolution of 120,000, AGC of 4 x 10^5^ ions, and max injection time of 50 ms, mass range of 350-1600 m/z; MS2 scans were acquired in the Orbitrap using the TurboTMT method with a resolution of 15,000, AGC of 1.25 x 10^5^, max injection time of 22 ms, HCD energy of 38%, isolation width of 0.4 Da, intensity threshold of 2.5 x 10^4^ and charges 2-6 for MS2 selection. Advanced Peak Determination, Monoisotopic Precursor selection (MIPS), and EASY-IC for internal calibration were enabled and dynamic exclusion was set to a count of 1 for 15 s.

#### Database search and post-processing analysis

All MS files were searched with Proteome Discoverer 2.4 using the Sequest node. Data was searched against the Uniprot Mouse database from Feb 2020 using a full tryptic digest, 2 max missed cleavages, minimum peptide length of 6 amino acids and maximum peptide length of 40 amino acids, an MS1 mass tolerance of 10 ppm, MS2 mass tolerance of 0.02 Da, variable oxidation on methionine (+15.995 Da) and fixed carbamidomethyl on cysteine (+57.021). Nat10-IP samples also included a fixed modification for TMTpro (+304.207) on lysine and peptide N-terminus and 4T1 cell lysates included variable modification for TMTpro (+304.207) on lysine and peptide N-terminus as well as acetylation on lysine (+42.011 Da). Percolator was used for FDR analysis and TMTpro reporter ions were quantified using the Reporter Ion Quantifier node and normalized on total peptide intensity of each channel.

### Immunofluorescence and confocal microscopy

Immunofluorescence analysis was carried out as described previously^8^. Briefly, cells were cultured on 4- or 8-well polymer coverslips (Ibidi) at seeding densities of 40,000 or 20,000 cells per well, respectively. After 24-48 h of incubation, cells underwent fixation with −20 °C methanol for 2 min and permeabilization with phosphate-buffered saline (PBS) containing 1% Triton X-100 for 1 min. Cells were then blocked with immunofluorescence buffer (1× PBS, 10 mg/ml bovine serum albumin (BSA), 0.02% SDS, and 0.1% Triton X-100) for 30 min. Subsequently, cells were incubated with primary antibodies diluted in immunofluorescence buffer overnight at 4 °C. After three washes with immunofluorescence buffer (5 min per wash), the cells were incubated with Alexa Fluor-conjugated secondary antibodies for 1 h at room temperature. Following three additional washes with immunofluorescence buffer, the cells were stained with 1 μg/ml DAPI for 10 min to label the nucleus. After three washes with PBS, slides were stored at 4 °C until subjected to confocal microscopy. In cases of drug treatment followed by super-resolution microscopy, cells were treated with 5-10 μM A-485, 10 μM BMH-21, and 20 ng/ml Actinomycin D for 24 h. Images were captured using either a Zeiss LSM 880 Airyscan super-resolution microscope (Airyscan detector, ×63 plan-apochromat NA 1.4 oil-immersion objective lens, and 0.05 μm X–Y pixel size), or a Nikon SoRa spinning disk super-resolution microscope (Yokogawa SoRa spinning disk unit, ×60 plan-apochromat NA 1.49 oil-immersion objective lens, Photometrics BSI sCMOS camera, and 0.027 μm X–Y pixel size). Airyscan images were processed using the Airyscan processing algorithm in the Zeiss ZEN Black (v.2.3) software, while Nikon SoRa images were deconvolved using a constrained iterative restoration algorithm in the Nikon NIS Elements (v5.11) software. Tetraspeck 0.2 μm beads (Invitrogen) were imaged with the same microscope parameters and used for channel alignment. Further image processing was carried out in Zeiss ZEN Blue V2 software. Primary antibodies for immunofluorescence were as follows: rabbit Nat10 (1:500, Abcam), mouse Nat10 (1:100, Proteintech), mouse H3.1/3.2 (1:1000, Active Motif), rabbit H3K9me3 (1:500, Abcam), rabbit p300 (1:250, Cell Signaling Technology), mouse H3K27ac (1:500, Active Motif), mouse H3K27me3 (1:1000, Active Motif), mouse SC35 (1:100, Abcam), mouse NPM1 (1:500, Abcam), mouse H4K20me3 (1:200, Active Motif). Secondary antibodies were used as follows: rabbit Alexa Fluor 568 (1:200, Invitrogen) and mouse Alexa Fluor 488 (1:200, Invitrogen).

### Proximity ligation assay

The proximity ligation assay was performed using the Duolink *In Situ* Orange starter kit mouse/rabbit (Millipore-Sigma) according to the manufacturer’s instructions. Briefly, 4T1 cells were grown on 15 well μ-slide angiogenesis coverslip (Ibidi). After 24h of incubation, cells were fixed in -20°C methanol for 2 min. After 3 washes with 1x PBS, cells were permeabilized with 1% Triton-X100 in PBS for 1 min. Cells were then blocked in Duolink blocking solution for 1 h at room temperature. Primary antibodies were diluted in Duolink dilution buffer at the ratios mentioned in the immunofluorescence protocol above. Incubation with primary antibody was performed overnight in a 4°C cold room. Cells were washed 2 times with 5% BSA in PBS for 10 min. PLUS, and MINUS secondary antibodies were diluted in antibody dilution buffer and added to cells for 1 h at 37°C in a humidified chamber. Cells were washed twice in 1x Buffer A for 5 min. Cells were then incubated with ligation mixture for 30 min at 37°C and subsequently washed twice with 1x Buffer A. Amplification mix was then added to cells and incubated for 100 min at 37°C. Cells were washed twice with 1x Buffer B for 10 min and then washed one time with 0.01x Buffer B for 1 min. Coverslips were then mounted with Prolong Gold Mounting Medium containing DAPI and imaged using a Zeiss LSM 880 Airyscan confocal microscope.

### Histone acetyl transferase (HAT) assay

Recombinant human p300 (full length), p300 catalytic subunit, CBP catalytic subunit, KAT7, NAT10, histone variants (H3.1, H3.2, H3.3, H4), and recombinant polynucleosomes were purchased from Active Motif. Recombinant human NAT10 derived from 293T cells was purchased from Origene Technologies. 5X HAT assay buffer (250 mM Tris-base, pH 8.0, 50% glycerol, 0.5 mM EDTA and 5 mM dithiothreitol) was purchased from Sigma. 250-500 μg of recombinant proteins were resuspended in HAT buffer supplemented with 0.75 mM acetyl CoA (Sigma) and 1 μl butyric acid (200 mM) (Sigma). Reaction volume was adjusted to 20 μl with H_2_O and incubated at 30°C for 30 min. Reactions were stopped with NuPAGE 4X LDS sample loading buffer (Thermo Fisher Scientific) and incubated at 95°C for 5 min. Samples were then loaded into an SDS-PAGE gel and western blotting protocol was followed as described above.

### (O-propargyl-puromycin) OPP assay

The OPP assay was performed using the Click-iT Plus OPP Alexa Fluor 488 Protein Synthesis Assay Kit (Thermo Fisher Scientific) according to the manufacturer’s instructions. Briefly, 4T1 cells were grown on glass coverslips placed in a 6-well tissue culture plate. After 48 h incubation, cells were treated with OPP reagent for 30 min. Cells were then fixed in 3.7% formaldehyde in PBS and permeabilized with 0.5% Triton-X100 in PBS. Subsequently, the Click-iT OPP detection procedure was performed according to the kit manual. Nuclei were then stained with 1 μg/ml DAPI followed by two washes with 1x PBS. Coverslips were mounted on glass slides using Prolong Glass Antifade Mounting Medium (Invitrogen) and imaged using a Zeiss LSM 780 confocal microscope.

### ac4C sequencing analysis of 18S rRNA

For ac4C sequencing of the helix 45 of 18S rRNA, total RNA was isolated using the standard Trizol method. Cells were directly lysed in TriPure lysis reagent (Sigma). After adding 200 μl chloroform, lysates were centrifuged at 14,000 rpm for 15min. 450 μl of the upper layer containing the RNA was separated in a fresh tube. Then 450 μl isopropanol was added to the solution and stored at -20°C for 1 h to precipitate RNAs. Precipitates were isolated by centrifugation at 14,000 rpm for 10 min, and then pellets were washed with 500 μl of 75% ethanol followed by centrifugation for 5 min. Pellets containing the total RNA were then resuspended in DEPC-treated H_2_O and treated with DNase I. 1 µg total RNA was treated with sodium cyanoborohydride (100 mM in H_2_O) or vehicle (H_2_O) in a final reaction volume of 100 μl. Reactions were initiated by the addition of 1 M HCl to a final concentration of 100 mM and incubated for 20 min at room temperature. Reactions were stopped by adding 30 μL of 1 M tris−HCl pH 8.0. the reaction volume was adjusted to 200 μl with H_2_O and RNA was purified via ethanol precipitation. The pelleted RNA was dried using a Speed-vac, resuspended in deioned H_2_O, and quantified using a Nanodrop 2000 spectrophotometer. For the reverse transcription reaction with the SuperScript III enzyme, ∼500 pg of treated RNA were incubated with 4.0 pmole of the helix 45 reverse primer (5’-TAATGATCCTTCCGCAGGTTCACCTAC-3’) in 1X Superscript III buffer at 65 °C for 5 min and transferred to ice for 1 min to facilitate annealing. After annealing, reverse transcriptions were performed by adding 200 units of the SuperScript III enzyme, 5 mM DTT, 25 units of RNasin, 500 μM dNTPs (5 mM GTP, 10 mM CTP, ATP, and TTP) and incubating at 55 °C for 60 min. Reactions were quenched by increasing the temperature to 70 °C for 15 min. The cDNA products from the reactions and controls were directly used in PCR. PCR reactions contained 2 μl cDNA in 50 μl PCR reaction with Phusion Hot Start Flex (New England Biolabs). Reaction conditions: 1X supplied HF buffer, 2.5 pmol each forward (5’-CGTCGCTACTACCGATTGGATGG-3’) and reverse (5’-TAATGATCCTTCCGCAGGTTCACCTAC-3’) primers, 200 μM each dNTP, 2 units of Phusion Hot Start enzyme, 2 μl template (Thermocycling conditions: 67 °C annealing, 34 cycles). PCR products were separated on a 2% agarose gel, stained with SYBR safe, and visualized on a UV transilluminator at 302 nm. Bands of the desired size were excised from the gel. DNA was extracted using a gel extraction kit (Zymo) and submitted for Sanger sequencing (GeneWiz) using 5 μM of the forward PCR primer. Processed sequencing traces were viewed using SnapGene software. Peak height for each base was measured, and the percent misincorporation was determined using the equation: “percent misincorporation = (peak intensity of T)/ (sum of C and T base peaks) X 100%”. Misincorporation values were determined by subtracting the background water control misincorporation levels from that of the corresponding reactions.

### ChIP-seq analysis

ChIP was performed using the ChIP-IT Express Enzymatic Chromatin Immunoprecipitation Kit from Active Motif, following the provided instructions. Initially, 4 × 10^6^ 4T1 cells were cultured on 15 cm tissue culture dishes for 48 h. Following incubation, cells were fixed with 1% formaldehyde for 5 min at room temperature and subsequently washed with ice-cold PBS. The formaldehyde cross-linking reaction was then stopped using a glycine stop-fix solution. Cells were collected by scraping and centrifuged at 2500 rpm for 10 min at 4 °C. Cells were lysed in ice-cold lysis buffer and homogenized using a Dounce homogenizer. Enzymatic shearing of chromatin was performed for 10 min at 37°C. The shearing process was stopped with 0.5 M EDTA, and the chromatin was isolated through centrifugation at 15,000 rpm for 10 min at 4°C. Each ChIP reaction was carried out using 25 μg of chromatin, along with 25 μl of Protein G magnetic beads and specific antibodies. The antibodies used included anti-rabbit H3K27ac (0.16 μg) (Cell Signaling Technology), anti-rabbit H3K27me3 (0.51 μg) (Cell Signaling Technology), anti-rabbit H4K20ac (5 μg) (Millipore-Sigma), and anti-rabbit p300 antibody (0.57 μg) (Cell Signaling Technology), anti-rabbit Brd4 (0.5 μg) (Cell Signaling Technology), anti-rabbit Nup210 (2 μg) (Bethyl Laboratories), anti-rabbit Rrp1b (1 μg) (Millipore-Sigma), anti-rabbit HA (5 μg) (Cell Signaling Technology), anti-rabbit V5 (1:40 dilution) (Cell Signaling Technology), and anti-rabbit SIPA (10 μg) (Abcam). Following an overnight incubation at 4 °C on a rotator, the beads were washed with ChIP buffers, and DNA was eluted using elution buffer. The eluted DNA was then subjected to reverse-cross linking and proteinase K treatment, with DNA quality assessed using an Agilent Bioanalyzer before ChIP-seq library preparation. The library was prepared using the SWIFT Accel-NGS 2s plus DNA Library Preparation Kit (Swift Biosciences), and the pooled samples were subsequently sequenced on the NovaSeq 6000 SP platform. ChIP-seq data peak calling was performed using the Sicer^30^ algorithm with a window size of 200 and gap size of 200. For the differential binding analysis, the Bioconductor package DiffBind was used after calling the peaks using the MACS2^31^ algorithm. ChIP-seq plot heatmap and plot profiles were created using deeptool2^32^.

### RNA isolation and quantitative reverse transcriptase real-time PCR (qRT-PCR)

For the total RNA isolation, TriPure Isolation Reagent (Sigma) was used to lyse cells directly on the cell culture plates. Following the addition of 200 μl of chloroform and centrifugation at 14,000 rpm. for 15 min at 4 °C, 450 μl of the upper aqueous layer, containing the RNA, was transferred to a new tube. To precipitate the RNA, 450 μl of isopropanol was added to each tube, followed by vortexing and incubation at −20 °C for 1 h. Subsequently, RNA purification was carried out using the RNeasy Mini Kit (Qiagen) with on-column DNase (Qiagen) digestion in accordance with the manufacturer’s instructions. 2 μg of total RNA was used for cDNA preparation using the iScript cDNA Synthesis Kit (Bio-Rad). A 1/10 dilution of the cDNA was then used for qRT-PCR analysis utilizing the SYBR Green PCR Master Mix (Applied Biosystems). The sequences of the primers employed are provided in the Supplementary Table 1.

### RNA-seq analysis

4T1 cells were grown in 6 cm tissue culture dishes, with a seeding density of 3 × 10^5^ cells per dish. Three replicate dishes were used for each condition. Following 48 h of incubation, total RNA was isolated using the RNeasy kit (Qiagen) according to the manufacturer’s instruction. To eliminate DNA contamination, on-column DNase treatment was applied using RNase-free DNase (Qiagen). For library preparation, the Stranded Total RNA Library Prep Kit (Illumina) was used, and the pooled samples were subjected sequencing on the NovaSeq 6000 system (Illumina). Analysis of differential gene expression from the RNA-seq data was conducted using the Partek Flow software. Gene Set Enrichment Analysis (GSEA) of the differentially expressed genes was carried out using the GSEA software (https://www.gsea-msigdb.org/gsea/index.jsp).

### 3D DNA Fluorescence *in Situ* Hybridization (FISH) analysis

3D DNA FISH analysis was performed with minor modifications using the protocol described elsewhere^33^. A BAC clone spanning the mouse Myc super-enhancer region (Clone ID: RP23-203H19) was purchased from Thermo Fisher Scientific. The BAC clone was purified using the NucleoBond Xtra BAC kit (Takara Bio). Alexa Fluor 488-labeled probe was generated through Nick translation system using the Chromatide Alexa Fluor 488-5-dUTP kit (Invitrogen). Labeled probe was precipitated and denatured at 85°C for 5 min, then pre-annealed at 37°C for 1 h and stored in a freezer until use. 4T1 cells were grown on μ-slide 15-well angiogenesis polymer coverslip (Ibidi) for 48 h. Cells were then fixed in 2% formaldehyde in PBS for 10 min at room temperature. After 3 rinses in 1x PBS, cells were permeabilized in ice-cold 0.4% Triton-X100 for 5 min, rinsed with PBS and then treated with 0.1 μg/μl RNase A (Sigma) for 1 h at 37°C. After 3 rinses in 1x PBS, cells were permeabilized with ice-cold 0.7% Triton-X100 + 0.1M HCl for 10 min. After 3 rinses in 1x PBS, cells were denatured in 1.9M HCl for 30 min at room temperature. After 3 rinses in PBS, cells were hybridized with labeled probes overnight at 37°C in a dark and humid chamber. Cells were washed 2 times with 2x SSC for 30 min in the dark (at 37°C and later at room temperature). Cells were then washed with 1x SSC for 30 min in the dark at room temperature. Nuclei were stained with 1 μg/ml DAPI for 10 min at room temperature. After 3 washes with 1x PBS, coverslips were mounted using Slow Fade Diamond Antifade Mountant (Invitrogen) and imaged using a Nikon SoRA spinning disk confocal microscope.

### Response to ECM stiffness assay

To investigate the impact of extracellular matrix (ECM) stiffness on tumor cells, we utilized polyacrylamide hydrogel-bound cell culture plates with varying elastic moduli to represent ECM stiffness. Specifically, 6-well plates with elastic moduli of 0.2 kPa (considered soft) was purchased from Matrigen. Regular 6-well tissue culture dishes were used to represent stiff matrices (elastic moduli is >GPa). Each well of the plate was treated with either 20 μg/ml fibronectin (Sigma) in PBS for 1 h at 37 °C or 50 μg/ml Type I Collagen (Gibco) overnight. 4T1 cells were then seeded onto either the soft or stiff matrix at a seeding density of 100,000 cells per well. After 48 h of incubation, cells were collected for protein isolation.

### High-parameter flow cytometric analysis of primary tumor-infiltrating leukocytes

4T1 WT and Nat10 KO-c43 primary tumors were harvested from BALB/c mice 34 days post-implantation. To obtain single cell suspensions, tumors were mechanically disrupted and then enzymatically digested for 1h at 37°C, with 1mg/ml of DNase (Roche, 10104159001) and 200U of collagenase IV (GIBCO, 17104-019) in 0.5% FCS RPMI 1640 media. The digested tissue was filtered through a 40 µm cell strainer followed by 3 min incubation with ACK lysing buffer at room temperature. The cell suspensions were then stained with LIVE/DEAD Fixable Dead Cell Stain kit (Invitrogen) in PBS followed by incubation with antibody cocktail prepared in Brilliant Stain Buffer (BD Biosciences, 563794). Fc receptors were blocked with anti-CD16/32 antibody (2.4G2, BioXcell). For biotinylated antibodies, cells were subsequently incubated with fluorochrome-conjugated streptavidin (BD Biosciences). The following antibodies were used: CD11b (M1/70), CD44 (IM7), CD45 (30-F11), CD86 (GL1), CD24 (M1/69), Ly6G (1A8), CD64 (X54-5/7.1), SiglecH (440C), PD-L1 (MIH5), MHCII (I-A/I-E) (M5/114.15.2), CD135 (A2f10.1), Ly6C (AL-21) from BD Biosciences; CD103 (2E7), Nkp46 (29.A1.4), ICAM1 (YN1/1.7.4), SiglecF (S17007L), CD206 (C068C2), F4/80 (BM8), CD127 (A7R34) from Biolegend; CD11c (N418), CD3e (145-2C11), TCRb (H57-597), TCRgd (GL-3) from ThermoFisher scientific.

Samples were acquired using a FACS Symphony A5 flow cytometer equipped with BD FACSDIVA software (BD Biosciences). Absolute cell counts were quantified using CountBright™ Absolute Counting Beads (Invitrogen) according to manufacturer’s instructions. Acquired data were analyzed as previously described using FlowJo^TM^ software (version 10.10) and R packages^34^. Tumor infiltrating leukocytes were selected following this gating strategy in FlowJo^TM^: debris were excluded using FSC-A versus SSC-A, doublets were excluded using FSC-H versus FSC-A, dead cells were removed using SSC-A versus Live/Dead positive cells, and tumor infiltrating leukocytes were selected using SSC-A versus CD45 positive cells. Asinh transformation, Phenograph clustering, T-Distributed Stochastic Neighbor Embedding (t-SNE), and Uniform Manifold Approximation Projection (UMAP) were ran combining all samples (15,000 cells/sample), using the R Spectre package^35^. Clusters were annotated based on the expression level of surface markers (Supplementary Table 2) and frequency of total leukocytes and each leukocyte population were extracted. Absolute number per mg of tumor was then calculated. ComplexHeatmap package^36^ was used to represent the level of expression of each marker of interest.

### Patient datasets analysis

Distant metastasis-free survival analysis of patients with high or low NAT10 expression was performed using the Km-plotter database (https://kmplot.com/analysis/). Analysis of NAT10 expression in the primary tumor and different metastatic sites of human breast cancer patients was performed from the datasets of the AURORA US Network submitted to the Gene Expression Omnibus (GSE193103) Database.

### Statistics and reproducibility

Experiments were conducted in duplicate or triplicate at minimum. In some cases of *in vivo* mouse studies, a single iteration was performed to minimize the use of animals. The robustness of key findings was confirmed by employing three different immunocompetent mice backgrounds in the metastasis assay, ensuring validation across diverse biological systems. Statistical analyses were performed using Graph Pad Prism 8 software. *In vivo* animal studies involved the calculation of p values using the Mann–Whitney U test for comparing two groups, and the outcomes were presented as mean ± standard deviation. In cases where more than two groups were compared, analysis of variance (ANOVA) with appropriate multiple comparison tests was performed. Statistical comparison of microscopy image analysis was performed either through the Mann– Whitney U test for the comparison of two groups or ANOVA with appropriate multiple comparison tests for the comparison of multiple groups. For the analysis of qRT-PCR results, p values were determined using multiple two-tailed, unpaired t-tests, and the results were reported as mean ± standard error of the mean (s.e.m).

### Microscopy image analysis and quantification

Nuclei stained for p300 and H3K9me3-marked/H3K27me3-marked heterochromatin or SC35 were segmented using the deep-learning Cellpose Python-package^37^. Nuclei were segmented using the pre-trained nuclei model, while p300 and other marked condensates were segmented using custom-trained models. Intensities and Mander’s colocalization coefficients were quantified using the scikit-image measure module^38^. p-values were calculated using Kruskal-Wallis ANOVA with Dunn’s multiple comparison test. Nucleolar segmentation was performed using intensity-based thresholding following median-filter denoising (kernel size = 2) based on NPM1 or NAT10 fluorescence intensity run in Python. Nucleolar area was calculated. Myc superenhancer distances to heterochromatin foci were quantified using Imaris image analysis software (Bitplane). Nuclei were segmented using the Surfaces module based on DAPI fluorescence intensity. Heterochromatin foci were segmented using the Surfaces module based on high-DAPI fluorescence intensity and Myc enhancer FISH signals were segmented using the Spots module. The shortest distance between each Myc FISH spot and heterochromatin foci surface was calculated. p-values were calculated using Kruskal-Wallis ANOVA with Dunn’s multiple comparison test.

### Data Availability

All the mass spectrometry data generated from this study have been deposited in the MassIVE (https://massive.ucsd.edu/ProteoSAFe/static/massive.jsp) database under the accession number: MSV00009365. RNA-seq and ChIP-seq data have been submitted to Gene Expression Omnibus (GEO) database under the accession numbers GSE250284 and GSE250428 respectively under the SuperSeries GSE250430.

## Result

### Depletion of NAT10 significantly decreases lung metastasis in mice

To determine whether NAT10 contributes to breast cancer metastasis, publicly available human breast cancer patient expression datasets were analyzed. Expression analysis of the primary tumor and metastases from the AURORA US Metastasis Project^39^ revealed that *NAT10* expression was significantly higher in breast cancer metastases compared to the primary tumor (Fig. 1a, b). Kaplan-Meier analysis of publicly available human breast cancer patient datasets revealed that higher expression of *NAT10* was associated with poor distant metastasis-free survival outcome in estrogen receptor positive (ER+), but not for triple negative breast cancer patients (Fig. 1c, d), consistent with a contributory role. To directly test the potential role of NAT10 in breast cancer metastasis, allograft-based spontaneous metastasis assays were performed. Orthotopic transplantation of *Nat10* shRNA knockdown (KD) mouse mammary tumor cells into BALB/cJ mice significantly decreased primary tumor weight (Fig. 1e, f) and lung metastases as compared with shRNA-Control (sh-Ctrl) 4T1 cells (Fig. 1g). To further verify the role of NAT10 in metastasis using an orthogonal method, we performed CRISPR/Cas9-mediated knockout (KO) of *Nat10* in 4T1 cells (Fig. 1h). However, as previously reported for HeLa cells^15^, we were not able to achieve complete knockout of *Nat10* in 4T1 cells as *Nat10* is an essential gene. Two independent *Nat10* KO clones (c19 and c43) showed inconsistent changes in primary tumor weight (Fig. 1i) but significantly decreased lung metastasis count (Fig. 1j). In addition, *Nat10* KD in the 6DT1 mouse mammary tumor cell line (Extended Data Fig. 1a) followed by orthotopic transplantation in FVB/NJ mice showed a similar trend of decreased lung metastasis without altering primary tumor weight (Extended Data Fig. 1b, c). In contrast, overexpression of *Nat10* in 4T1 significantly increased primary tumor weight (Extended Data Fig. 1d, e) and caused a trend toward increased lung metastasis (Extended Data Fig. 1f). Similar findings were observed upon overexpression of *Nat10* in 6DT1 cells, though primary tumor weight was not altered (Extended Data Fig. 1g, h), consistent with a role for NAT10 in metastatic disease. To further verify the role of NAT10 in metastasis, C57BL/6 mice carrying an H537A inactivating point mutation in the *Nat10* acetyl transferase domain (Meier et al., manuscript in preparation) were crossed with FVB/NJ male mice of the highly metastatic PyMT mammary tumor model (Fig. 1k) and tumor and metastasis phenotypes in the F1 female progeny were assessed. Consistent with the orthotopic transplantation models, mice carrying one copy of the mutant Nat10 allele had significantly decreased lung metastases (Fig. 1l, m) and metastatic incidence (Fig. 1n), confirming a role of NAT10 in metastatic progression and specifically implicating the NAT10 acetyl transferase function in this phenotype.

**Fig. 1.**
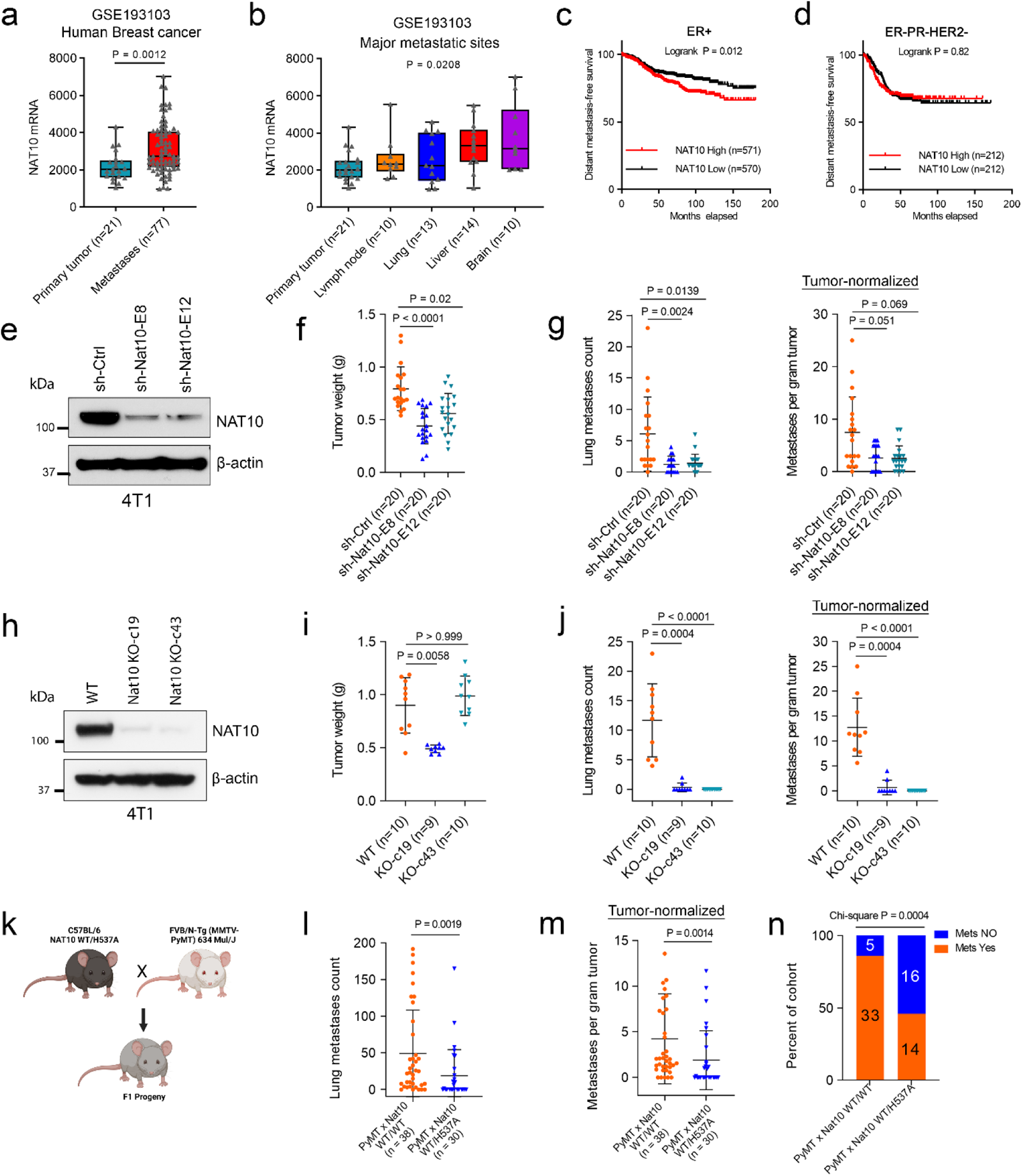
NAT10 depletion decreases lung metastasis in mice. (a) Level of NAT10 mRNA in the primary tumor and metastases of human breast cancer patient. Mann-Whitney U test, error bar represents median with interquartile range. (b) Level of *NAT10* mRNA in different metastatic sites of human breast cancer patient, error bar represents median with interquartile range. Kaplan-Meier survival curve showing the distant metastasis-free survival outcome of human (c) ER+ and (d) triple-negative (ER-/PR-/HER2-) breast cancer patients with high or low *NAT10* expression. Logrank test. (e) Western blot showing the level of NAT10 protein in shRNA knockdown 4T1 cells. (f) Tumor weight of mice injected with *Nat10* shRNA knockdown 4T1 cells. Kruskal Wallis ANOVA, mean ± S.D. p-values from two independent experiments were combined with Fisher’s method. (g) Lung metastases count (left) and tumor-normalized metastases count (right) in *Nat10* shRNA knockdown 4T1 cells. Kruskal-Wallis ANOVA with Dunn’s multiple comparison test, mean ± s.d. p-values from two independent experiments were combined with Fisher’s method. (h) Western blot showing the level of NAT10 in *Nat10* KO 4T1 cells. (i) Tumor weight of mice injected with *Nat10* KO 4T1 cells, Kruskal-Wallis ANOVA with Dunn’s multiple comparison test, mean ± s.d. (j) Lung metastases count (left) and tumor-normalized metastases count in *Nat10* KO 4T1 cells. Kruskal-Wallis ANOVA with Dunn’s multiple comparison test, mean ± s.d. (k) Breeding scheme for crossing *Nat10* mutant (H537A) mice with PyMT mice. (l) Lung metastases count and (m) tumor-normalized metastases count in WT and *Nat10* mutant F1 progeny. Mann-Whitney U test, mean ± s.d. (n) Incidence of lung metastasis in WT and *Nat10* mutant F1 progeny. Chi-square test.

### Protein translational control pathway is active in NAT10 depleted mammary tumor cells

NAT10 has previously been described as an RNA acetyl transferase, catalyzing cytidine acetylation (ac4C) of 18S RNA and tRNAs^16^. Accordingly, we performed ac4C analysis of helix 45 of 18S rRNA (Extended Data Fig. 2a) from *Nat10* KO 4T1 cells to functionally assess our knockout efficiency. As expected, the ac4C level of 18S rRNA was significantly reduced in the NAT10 depleted cells (Extended Data Fig. 2b). Recently, acetylation of mRNAs by NAT10 has been implicated in increased protein translation efficiency^15^. Therefore, it is conceivable that reduction of Nat10 activity, either by the inactivating point mutation in the mouse model or by knocking out/down in cell lines, would affect protein translational capacity. Unexpectedly, western blot analysis revealed a slight increase in both total p70S6 kinase and phospho-p70S6 kinase levels in *Nat10* KO cells, indicative of an activation of translational control pathways (Extended Data Fig. 2c). Furthermore, *Nat10* KO did not induce translational stress in this cell line as confirmed by unaltered level of phospho-EIF2α. Consistent with this, O-propargyl-puromycin (OPP) assays on NAT10-depleted cells showed an observable increase in nascent protein synthesis (Extended Data Fig. 2d). These results suggest that the mechanism by which NAT10 facilitates metastasis in our system is unlikely to be via modulation of protein translation.

### NAT10 depletion phenocopies disruption of the mechanosensitive NUP210-dependent metastasis pathway

Unexpectedly, NAT10-depleted cells exhibited a significant adhesion defect on both glass and plastic matrices (Fig. 2a). We recently observed a similar adhesion defect in tumor cells depleted of NUP210, a nuclear pore-associated mechanosensitive and metastasis-promoting protein^8^. Since NUP210 and NAT10 interact with a similar cadre of metastasis susceptibility proteins, including BRD4-SF, SIPA1 and RRP1B^13^, it follows that NAT10 might function in the putative NUP210-dependent mechanosensitive pathway in metastasis (Fig. 2b). Coimmunoprecipitation (Co-IP) analysis revealed that NUP210 strongly pulled down NAT10, but only weakly associated with SIPA1 (Fig. 2c). Flag-tagged NAT10 interacted with both the long- and two short isoforms of endogenous BRD4 in 4T1 cells (Fig. 2d). Reciprocal Co-IP in human 293FT cells revealed that NAT10 interacts with both isoforms of BRD4, but the interaction with BRD-SF might be stronger (Fig. 2e, f). Consistent with the Co-IP data, proximity ligation assays further demonstrated an interaction of NUP210 with both NAT10 and SIPA1 (Fig. 2g), suggesting that NAT10 was a member of the NUP210-associated protein complex at the nuclear pore. Interestingly, depletion of NAT10 was associated with a significant decrease of NUP210 and integrin β1 protein levels (Fig. 2h). NAT10 depletion also significantly decreased the expression of NUP210-regulated mechanosensitive, inflammatory response genes (Fig. 2i). As expected, similar to NUP210-depleted cells, immunofluorescence analysis showed redistribution of histone H3.1/3.2 to H3K9me3-enriched heterochromatin foci after NAT10 depletion (Fig. 2j). Moreover, like NUP210 upregulation in higher stiffness matrices, protein levels of NAT10, RRP1B, SIPA1 and BRD4 isoforms were upregulated on plastic tissue culture plates (stiffness > GPa) compared to soft (stiffness 0.2 kPa) matrices (Fig. 2k). Taken together, these data are consistent with NAT10 functioning together with NUP210 at the nuclear pore as part of the mechanosensitive pathway that promotes metastasis.

**Fig. 2.**
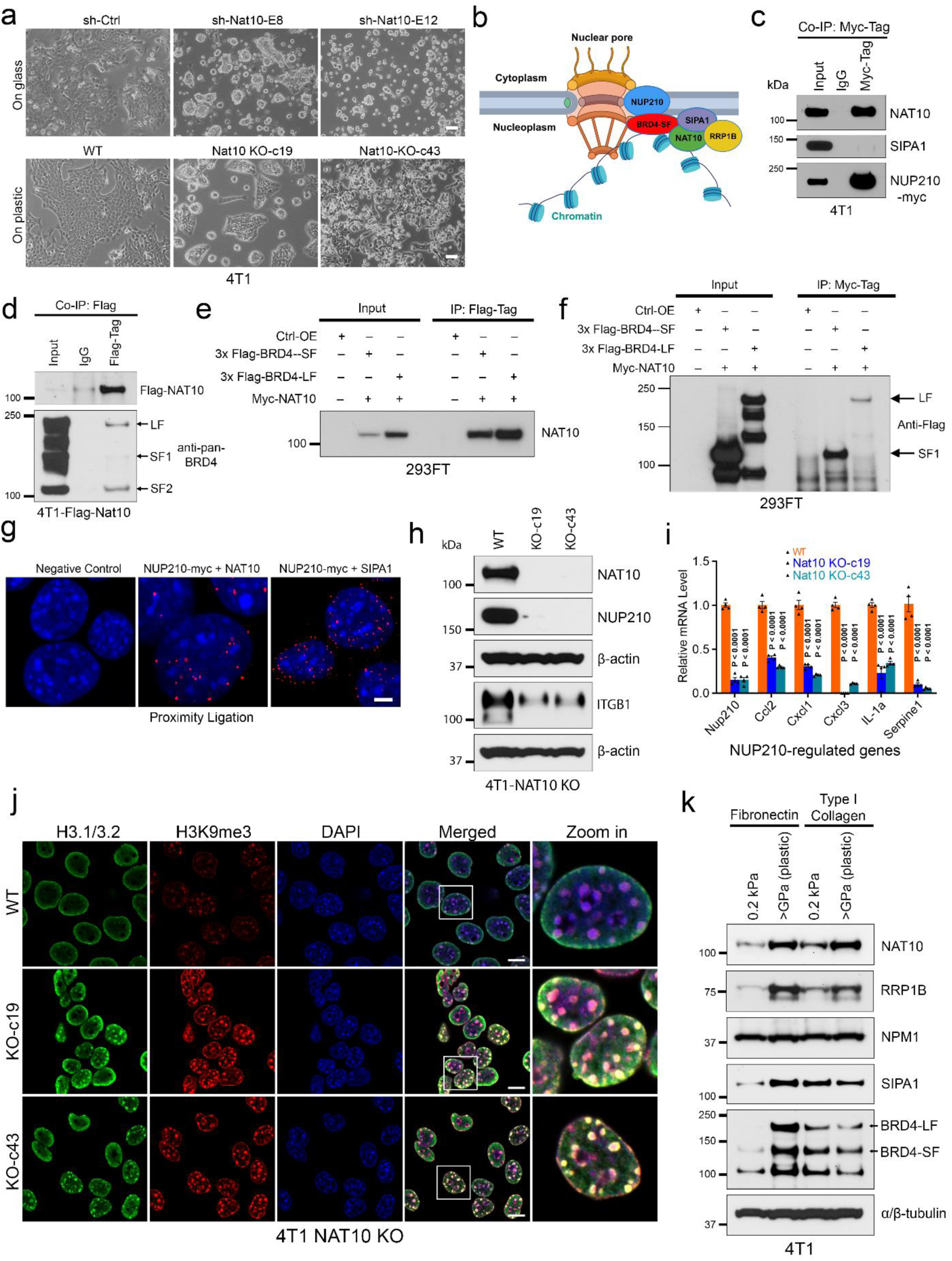
NAT10 interacts with a metastasis-associated, nuclear pore-bound mechanosensitive protein complex. (a) Morphology of *Nat10* shRNA knockdown (top) and *Nat10* KO 4T1 cells grown on different plates (bottom). Scale bar = 10 μm. (b) Hypothetical model of the interaction between NAT10 and the NUP210-bound mechanosensitive protein complex at the nuclear pore. (c) Co-IP showing the interaction of NAT10 with Myc-tagged NUP210 and SIPA1 in 4T1 cells. (d) Co-IP showing the interaction of Flag-tagged NAT10 and BRD4 isoforms in 4T1 cells. (e) and (f) Reciprocal Co-IP showing the interaction of Myc-tagged NAT10 and Flag-tagged BRD4 isoforms in human 293FT cells. (g) Proximity ligation assay showing the interactions (red dots) of NUP210 with NAT10 and SIPA1. Scale bar = 5 μm. (h) Western blot showing the level of NUP210 and ITGB1 protein in *Nat10* KO 4T1 cells. (i) qRT-PCR showing the level of NUP210-dependent mechanosensitive, inflammatory response genes in *Nat10* KO 4T1 cells, multiple t-test, mean ± s.e.m. (j) Immunofluorescence showing the distribution of histone H3.1/3.2 and H3K9me3 heterochromatin markers in *Nat10* KO 4T1 cells. Scale bar = 10 μm. (k) Western blot showing the levels of NAT10 and associated mechanosensitive proteins in 4T1 cells grown on plates with soft (0.2kPa) and stiff (plastic dish, stiffness > GPa) matrices coated with either fibronectin or type I collagen.

### Depletion of NAT10 leads to decreased histone acetylation

In addition to the known role as an ac4C acetylase for mRNAs^15^, rRNAs and tRNAs^16^, NAT10 has been reported to acetylate proteins such as UBF^17^, tubulin^40^, p53^18^, and MORC2^19^. Moreover, NAT10 itself is acetylated, which is thought to be critical for UBF acetylation and 18S rRNA transcription^41^. Since NAT10 depletion was associated with redistribution of histone H3.1/3.2 and transcriptional suppression of NUP210-target genes, we hypothesized that NAT10 might be acetylating chromatin-associated proteins and affecting gene transcription. To identify other potential acetylation targets of NAT10 that might be associated with transcriptional regulation in metastasis, an unbiased mass spectrometry-based approach was performed. Immunoprecipitation-mass spectrometry of nuclear proteins revealed that 361 proteins potentially interact with NAT10 in 4T1 cells (Fig. 3a, b). Gene Set Enrichment Analysis (GSEA) analysis of these proteins identified ‘MYC targets’ and ‘positive regulation of transcription by RNA polymerase II’ as top enriched pathways. (Fig. 3b, c). To identify potential direct acetylation targets of NAT10, proteome-normalized acetylated lysine peptide enrichment was performed in WT and *Nat10* KO 4T1 cells. Several histones were identified among the top downregulated acetylated proteins in *Nat10* KO 4T1 cells (Fig. 3d). Overlap analysis of the two proteomics datasets revealed that nine NAT10-interacting proteins had reduced acetylation NAT10 depleted cells (Fig. 3e), including multiple histone proteins, consistent with changes in the H3.1/3.2-associated chromatin structure observed in the immunofluorescence analyses. To further investigate whether histones are substrates of NAT10 in 4T1 cells, pan-acetylated lysine (K) immunoprecipitation followed by western blot analysis was performed in WT and *Nat10* KO 4T1 cells. Decreased acetylation of three major histone variants (H3.1/3.2, H3.3 and H4) without alteration of their total concentration was found in *Nat10* KO 4T1 cells (Fig. 3f). Western blot analysis further confirmed that NAT10 depletion was associated with decreased histone H3 and H4 acetylation (Fig. 3g).

**Fig. 3.**
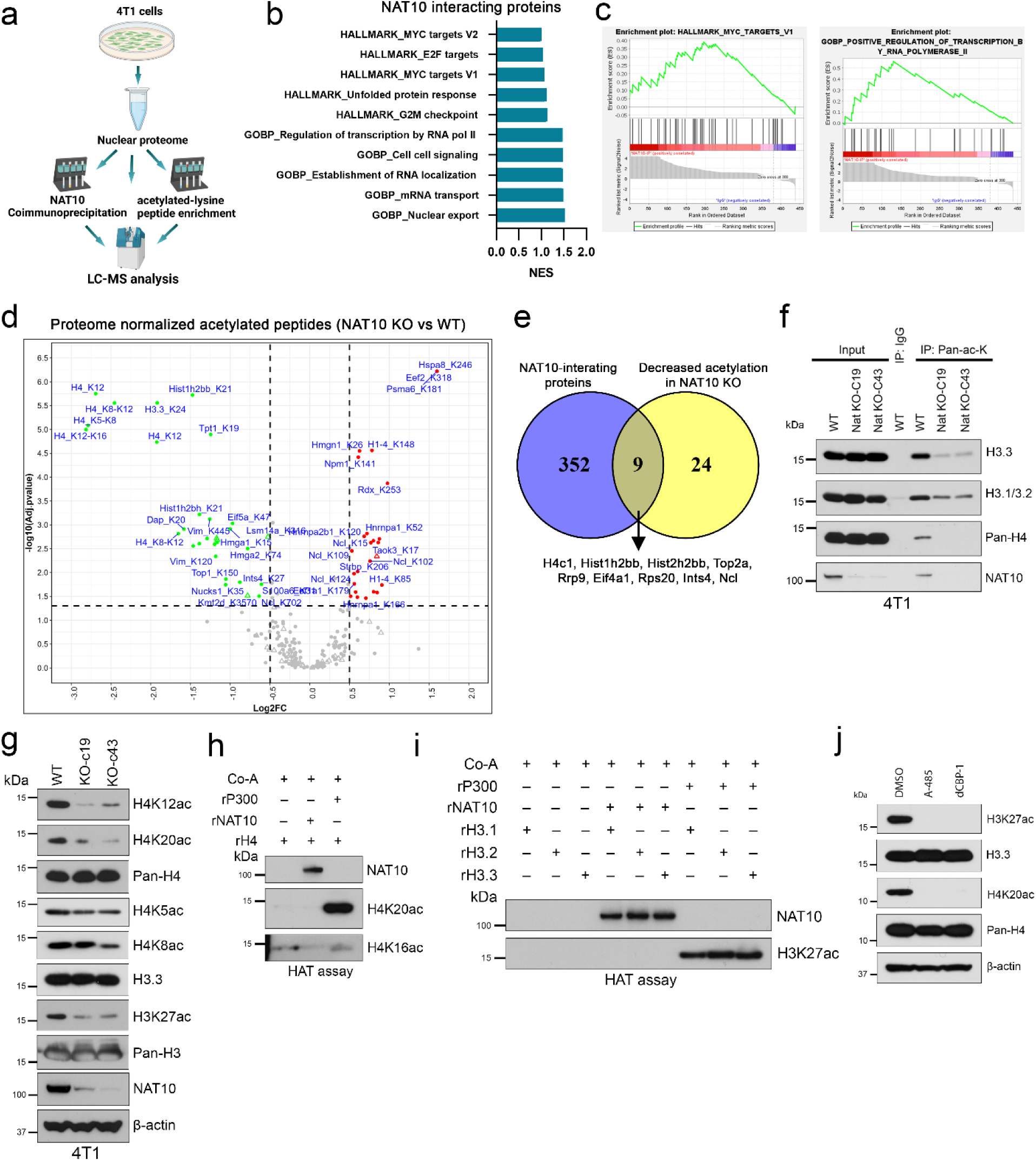
NAT10 depletion decreases global histone acetylation. (a) Scheme used for combined LC-MS-based identification of NAT10 interaction partners and acetylated lysine enrichment analysis in *Nat10* KO 4T1 cells. (b) GSEA analysis of NAT10-interacting proteins identified through Co-IP followed by mass spectrometry in 4T1 cells. (c) Key GSEA pathways enriched among NAT10-interacting proteins. (d) Volcano plot showing the nuclear proteome-normalized acetylated peptides in *Nat10* KO 4T1 cells. (e) Venn diagram showing the overlap of NAT10 interaction partners and proteins with decreased acetylation in *Nat10* KO 4T1 cells. (f) Co-IP showing decreased pull down of acetylated histones in *Nat10* KO 4T1 cells. (g) Western blot showing the levels of different histone modifications in *Nat10* KO 4T1 cells. (h) *In vitro* histone acetyltransferase (HAT) assay showing the level of histone H4 acetylation in the presence of recombinant human NAT10 and p300. (i) *In vitro* HAT assay showing the acetylation level of histone H3 variants in the presence of recombinant human NAT10 and p300. (j) Western blot showing the level of H3K27ac and H4K20ac in 4T1 cells treated with either a p300/CBP inhibitor (A-485) or a p300/CBP PROTAC degrader (dCBP-1).

Nat10 is an essential gene, and its most well-characterized targets are tRNA and rRNA. Therefore, we took an open posture as to whether the observed changes in histone acetylation may be direct or indirect. To test whether NAT10 can directly acetylate histone H3 and H4, *in vitro* histone acetyl transferase activity (HAT) assays were performed using recombinant human NAT10 protein. However, the *in vitro* HAT assay revealed that p300, not NAT10, was able to acetylate nucleosome-free H4 and H3 variants (Fig. 3h, i). To confirm whether p300 is involved in H3K27ac and H4K20ac modification in the cellular context, 4T1 cells were treated with either a p300/CBP inhibitor (A-485) or p300/CBP PROTAC degrader (dCBP-1). Both H3K27ac and H4K20ac modifications were drastically reduced in p300 inhibitor-treated cells, suggesting that p300 is the main HAT for these modifications (Fig. 3j). To test whether NAT10 can acetylate histones in the nucleosome context, we performed the HAT assay using recombinant polynucleosomes and two different recombinant human NAT10 proteins (derived from either human 293T cells or bacuolovirus). Again, the p300 catalytic subunit, but not NAT10 could acetylate histones (Extended Data Fig. 3a). Since p300 is itself regulated by acetylation, we tested whether NAT10 could indirectly affect histone acetylation by regulation of p300 activity or it’s paralog CREB-binding protein (CBP). However, addition of NAT10 to HAT assays did not alter the acetylation levels of the p300 or CBP catalytic subunits (Extended Data Fig. 3b, c). Additionally, NAT10 did not alter the acetylation levels of another histone acetyltransferase, KAT7/HBO1 (Extended Data Fig. 3d). While we cannot completely dismiss the protein acetyltransferase activity of NAT10, the weight of our evidence indicates that NAT10 is not capable of directly acetylating proteins, histones, or histone acetytransferase p300. This led us to explore whether NAT10 might indirectly regulate protein acetyltransferase activity.

### NAT10 depletion promotes p300 upregulation and mislocalization

Since NAT10 depletion resulted in decreased acetylation of p300 targets, western blot analysis was performed to assess p300 levels in KO cells. Unexpectedly, both p300 levels and acetylated p300/CBP (ac-p300 K1499/ac-CBP K1535) were significantly upregulated in *Nat10* KO 4T1 cells (Fig. 4a). In contrast, re-expression of NAT10 in KO cells downregulated the p300 protein level (Fig. 4b). Similar results were also observed in 6DT1 cells with shRNA-mediated knockdown of *Nat10* (Extended Data Fig. 4a) suggesting that the p300 dysregulation was not an off-target effect. Finally, to eliminate the possibility of an artifact due to stable depletion of NAT10 through CRISPR/Cas9 or shRNAs, NAT10 depletion through dTAG degron tagging^29^ in human 293FT cells was performed. Acute depletion of NAT10 in 293FT cells also resulted in upregulation of p300 (Extended Data Fig. 4b). These results suggest that NAT10 is an obligatory co-factor for p300-associated transcriptional regulatory complex and cells increase the level of p300 to compensate for the loss of p300 activity in absence of NAT10.

**Fig. 4.**
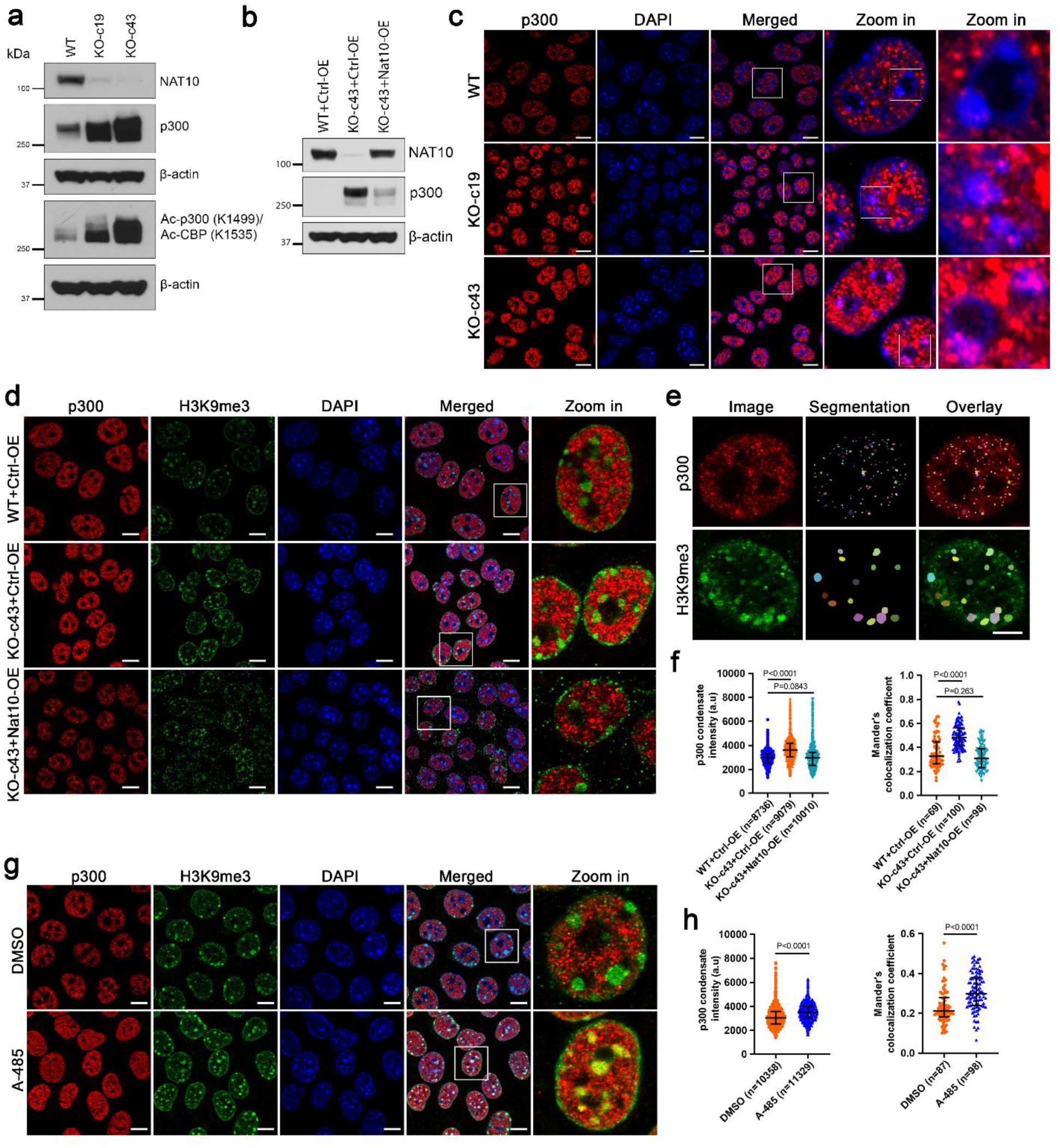
NAT10 depletion leads to mislocalization of p300 into heterochromatin zones. (a) Western blot showing the levels of p300 and acetylated p300 in *Nat10* KO 4T1 cells. (b) Western blot showing the level of p300 in *Nat10* KO and rescued (by overexpression of *Nat10*) 4T1 cells. (c) Immunofluorescence showing the distribution of p300 in *Nat10* KO 4T1 cells, Scalebar = 10 μm. (d) Immunofluorescence showing the distribution of p300 and H3K9me3 in *Nat10* KO and rescued 4T1 cells, Scalebar = 10 μm. (e) Segmentation criteria of the p300 and H3K9me3 signals for quantification. Scalebar = 5 μm. (f) Quantification of p300 condensate intensity in *Nat10* KO and rescued 4T1 cells (left), Mander’s colocalization coefficient of p300 localization in the H3K9me3 area (right). Kruskal-Wallis ANOVA with Dunn’s multiple comparison test, median with interquartile range. (g) Immunofluorescence showing the distribution of p300 and H3K9me3 in A-485 treated 4T1 cells, Scalebar = 10 μm. (h) Quantification of p300 condensate intensity in A-485 treated 4T1 cells (left), Mander’s colocalization coefficient of p300 localization in the H3K9me3 area (right). Kruskal-Wallis ANOVA with Dunn’s multiple comparison test, median with interquartile range.

To determine whether the distribution of p300 is altered in *Nat10* KO cells, immunofluorescence analysis was performed. In *Nat10* KO cells, p300 was found to be mislocalized in heterochromatin regions (strong DAPI signal area) (Fig. 4c). To further confirm the association of p300 with heterochromatin, immunostaining with the heterochromatin marker H3K9me3 was performed. Quantification of p300 condensate signals in *Nat10* KO and rescued cells revealed that p300 condensate intensity was significantly increased and colocalized with H3K9me3 marks in *Nat10* KO cells which was rescued after re-expression of *Nat10* in KO cells (Fig. 4d-f). To assess whether the mislocalization of p300 is associated with loss of p300 activity, WT cells were treated with a p300 inhibitor A-485 or a p300/CBP PROTAC degrader (dCBP-1). Western blot analysis demonstrated an increase in p300 levels in A-485 treated cells, which was accompanied by loss of H3K27ac (Extended Data Fig. 4c). In addition, consistent with the Nat10 KO cells, immunofluorescence analysis revealed that A-485 treatment significantly increased nuclear p300 intensity and caused re-localization of p300 into H3K9me3 heterochromatin regions (Fig. 4g, h).

p300 transcriptional activity is negatively regulated by phosphorylation at serine 89^42^. To test whether increased levels of p300 were due to increased S89 phosphorylation in *Nat10* KO cells, western blot analysis was performed. p-p300 (S89) was increased in *Nat10* KO cells, consistent with the observed p300 inactivation. However, re-expression of *Nat10* in the KO cells did not significantly rescue the p-S89 levels (Extended Data Fig. 4d) suggesting that this inactivating phosphorylation event is not a direct effect of NAT10 depletion and may not play a role in the mislocalization phenotype.

A recent report suggested that the HAT activity of p300 can be reduced due to the occlusion of active sites in phase-separated p300 condensates^43^. A large proportion of these p300 condensates can also colocalize with H3K27me3 facultative heterochromatin marks. We therefore tested whether p300 condensates colocalized with H3K27me3 in the *Nat10* KO 4T1 cells. Indeed, H3K27me3 was significantly increased and colocalization of p300 condensates with H3K27me3 was significantly increased in *Nat10* KO cells (Extended Data Fig. 4e, f) implying that p300 might lose its HAT activity due to the occlusion of its active site in the heterochromatin region. To test whether p300 is also mislocalized in inter-chromatin regions, staining with SC35, an inter-chromatin granule nuclear speckle marker, was performed. Again, colocalization of p300 with SC35 was significantly increased in *Nat10* KO cells (Extended Data Fig. 4g, h), indicating that NAT10 depletion can also lead to p300 mislocalization in the inter-chromatin space. Taken together, these results suggest that NAT10 is required for proper p300 chromatin acetylation and localization.

### NAT10 depletion disrupts enhancer organization

Since p300 is essential for histone acetylation dependent enhancer activation^44^, enhancer organization was examined in *Nat10* KO cells. ChIP-seq analysis revealed that NAT10 depletion decreased the overall enrichment of H3K27ac and H4K20ac enhancer marks as well as p300 in gene bodies (Fig. 5a, b) but not at the transcriptional start sites (TSS; Extended Data Fig. 5a, b). There was no significant change in the enrichment of H3K27me3 facultative heterochromatin marks. Differential ChIP enrichment analysis of enhancer marks identified 9928 downregulated- and 6161 upregulated H3K27ac peaks in *Nat10* KO 4T1 cells (Fig. 5c). Similarly, 7098 downregulated- and 12866 upregulated H4K20ac peaks were identified *Nat10* KO cells. Analysis of the peak distribution within the genes revealed that 1999 genes had downregulated (1.5-fold) and 929 genes had upregulated (1.5-fold) H3K27ac peaks enrichment in *Nat10* KO cells (Fig. 5d). Gene ontology analysis revealed that genes with downregulated H3K27ac enhancer marks were mainly cell adhesion and migration-related genes. Consistent with their decreased expression, decreased enrichment of H3K27ac and H4K20ac marks was found in the cell adhesion genes *Nup210* and *Itgb1* (Extended Data Fig. 5c). In contrast, genes with upregulated H3K27ac peaks were mainly developmental, differentiation-related genes. Similar observations were made for the gene ontology analysis of differential H4K20ac peaks (Extended Data Fig. 5d).

**Fig. 5.**
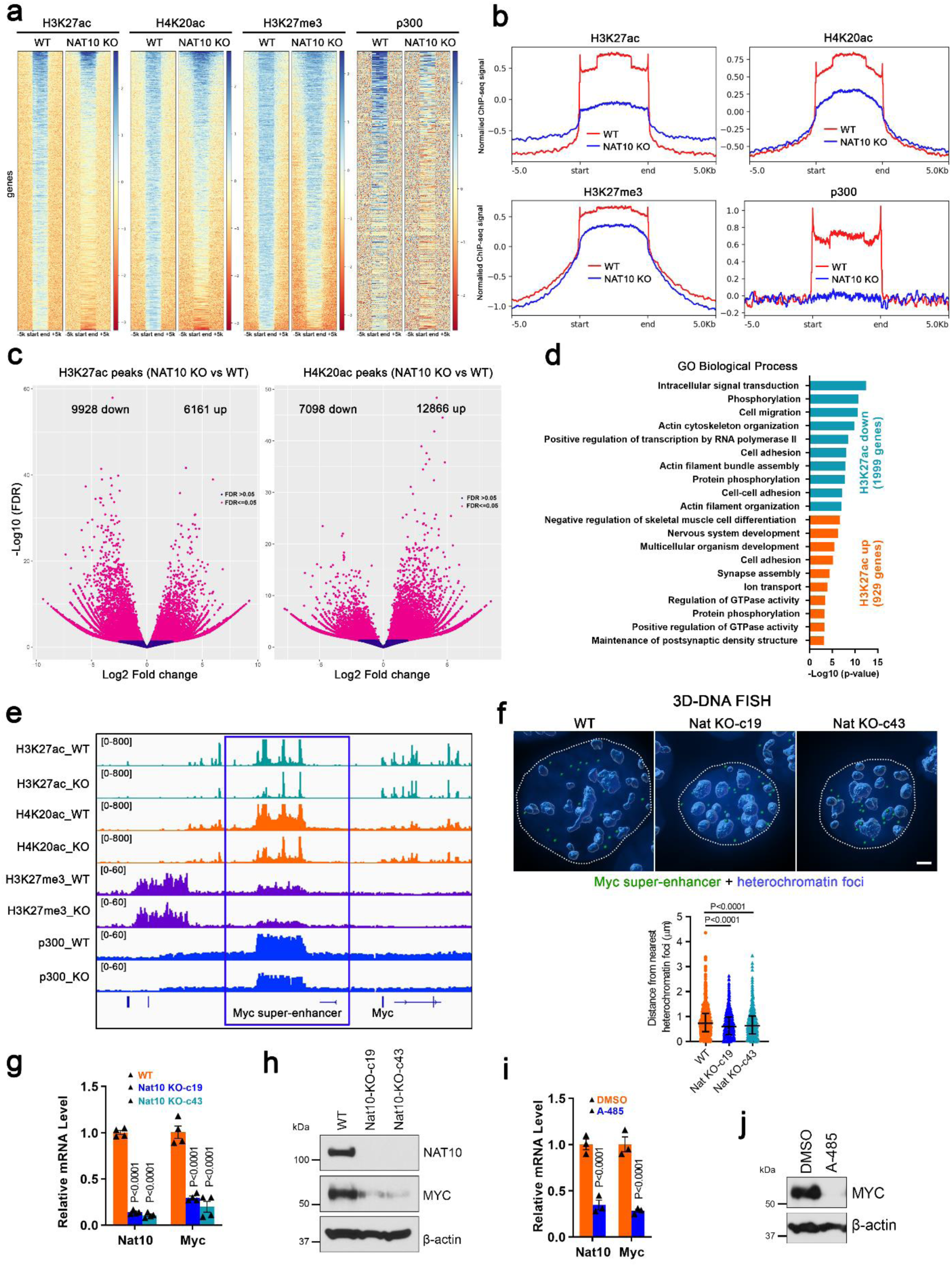
NAT10 depletion disrupts enhancer organization. (a) ChIP-seq heatmap showing the peak distribution of H3K27ac, H4K20ac, H3K27me3 and p300 within gene bodies of *Nat10* KO 4T1 cells. (b) Normalized ChIP enrichment profile of H3K27ac, H4K20ac, H3K27me3 and p300 in *Nat10* KO 4T1 cells. (c) Differential binding analysis of H3K27ac and H4K20ac peaks in *Nat10* KO 4T1 cells. (d) Gene Ontology analysis of down- and upregulated enrichment of H3K27ac peaks within gene bodies in *Nat10* KO 4T1 cells. (e) Integrative genomic viewer (IGV) plot showing the differential enrichment of H3K27ac, H4K20ac, H3K27me3, and p300 within the *Myc* super-enhancer region in *Nat10* KO 4T1 cells. (f) 3D DNA FISH analysis showing the distribution of *Myc* super-enhancer region within the nuclei of *Nat10* KO 4T1 cells (top), Scalebar = 2 μm; quantification of the distance of FISH signals from the nearest heterochromatin foci (bottom). Kruskal-Wallis ANOVA with Dunn’s multiple comparison test, median with interquartile range. (g) qRT-PCR showing the mRNA level of *Nat10* and *Myc* in *Nat10* KO 4T1 cells, multiple t-test, mean ± s.e.m. (h) Western blot showing the protein level of MYC in *Nat10* KO 4T1 cells. (i) qRT-PCR showing the mRNA level of *Nat10* and *Myc* in A-485 treated cells, multiple t-test, mean ± s.e.m. (j) Western blot showing the level of MYC in A-485-treated 4T1 cells.

NAT10 depletion also led to decreased enhancer marks and decreased p300 binding in *Myc* super-enhancer region (Fig. 5e). To verify whether NAT10 depletion alters overall enhancer organization, we performed 3D DNA FISH analysis to visualize the intra-nuclear localization of the *Myc* super-enhancer region. *Myc* super-enhancer loci were highly amplified in aneuploid 4T1 cells (Fig. 5f). Quantification of the distance between the Myc super-enhancer loci and the nearest heterochromatin foci revealed that Myc super-enhancer loci became closer to the heterochromatin foci in *Nat10* KO cells (Fig. 5f). Decreased *Myc* mRNA (Fig. 5g) and protein expression was also observed in *Nat10* KO cells (Fig. 5h). Treatment of WT 4T1 cells with a p300 inhibitor A-485 treatment also downregulated *Myc* mRNA (Fig. 5i) and protein expression (Fig. 5j). These results are therefore consistent with the depletion of NAT10 disrupting p300 enhancer localization and altering enhancer organization.

### NAT10 interacts with p300 at enhancers

To assess whether NAT10 mediates its effect on histone acetylation via interaction with p300, Co-IP with either endogenous or Flag-tagged NAT10 in 4T1 cells was performed. Co-IP revealed that both Flag-tagged and endogenous NAT10 physically interacts with p300 (Fig. 6a). Immunofluorescence analysis revealed that nucleoplasmic and perinucleolar NAT10 colocalized with p300 and H3K27ac enhancer marks (Fig. 6b). To further confirm the association of NAT10 with enhancers, proximity ligation assays in 4T1 cells were performed. NAT10 was found to strongly interact with H3K27ac and H4K20ac enhancer marks (Fig. 6c). 3D reconstruction of the images revealed that many of these interaction sites were in the nucleoplasm, near heterochromatin foci (dark blue foci) and the nucleolar periphery (dark zone surrounded by heterochromatin foci) (Fig. 6d). To further validate the interaction of NAT10 with enhancers, we performed ChIP-seq analysis of NAT10, H3K27ac and H4K20ac. Overlaying the enrichment signal of NAT10 with these enhancer marks revealed that NAT10 was diffusely enriched in the H3K27ac and H4K20ac enhancer mark regions (Fig. 6e). Interestingly, ChIP-seq analysis of other components of nuclear pore-bound mechanosensitive protein complex revealed that these proteins were enriched in the *Myc* super-enhancer region (Fig. 6f). These results suggested that NAT10 interacts with p300 along with other nuclear pore-bound mechanosensitive proteins at enhancer regions to facilitate histone acetylation.

**Fig. 6.**
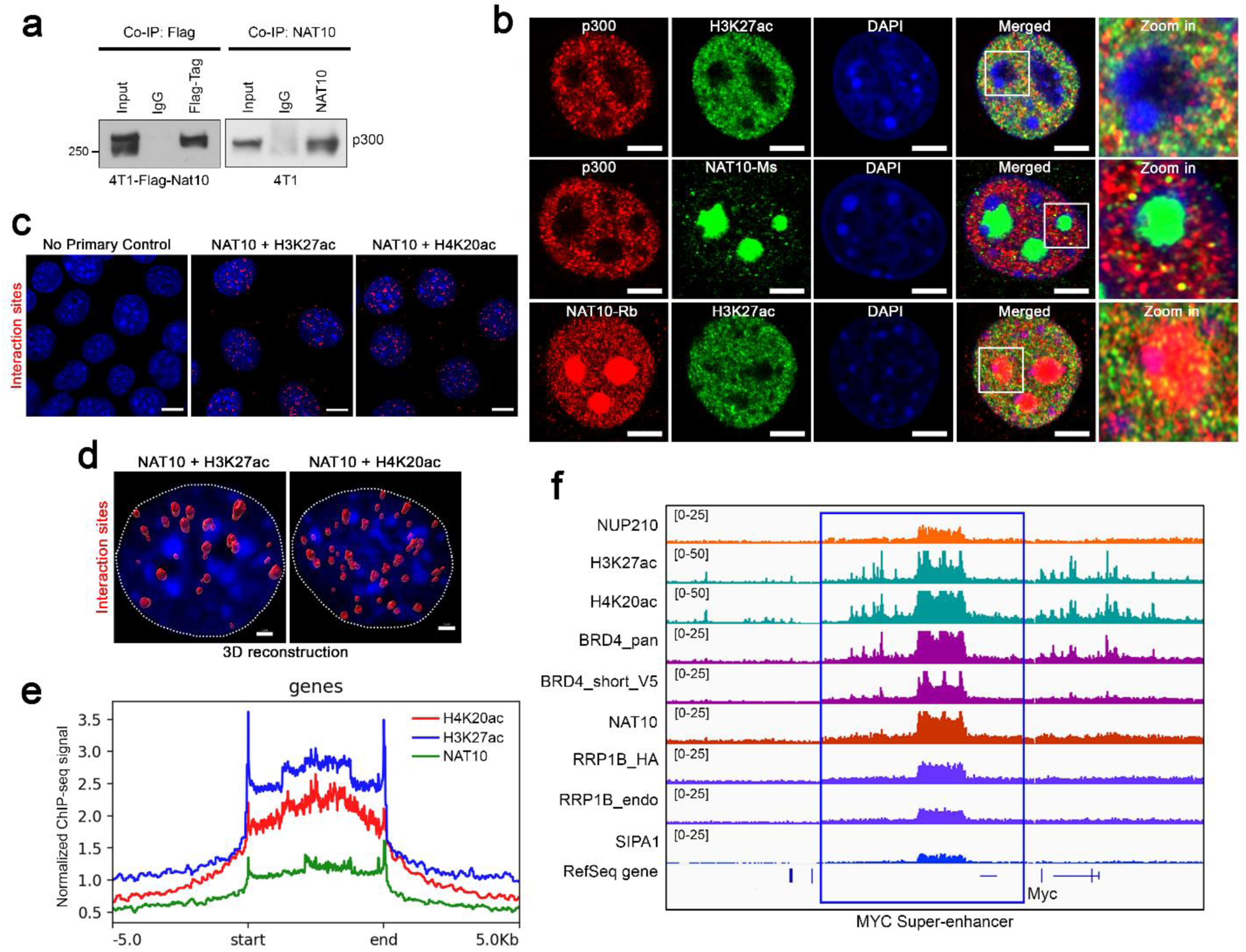
NAT10 interacts with p300-associated enhancers. (a) Co-IP showing the interaction of Flag-tagged and endogenous NAT10 with p300 in 4T1 cells. (b) Immunofluorescence showing the colocalization of NAT10 with p300 and enhancer mark H3K27ac, Scalebar = 5 μm. NAT10-Ms, mouse NAT10 antibody; NAT10-Rb, rabbit NAT10 antibody. (c) Proximity ligation assay showing the interaction of NAT10 with H3K27ac and H4K20ac marks in 4T1 cells, Scalebar = 10 μm. (d) 3D reconstruction showing NAT10 interactions with H3K27ac and H4K20ac within the nucleus of 4T1 cells, Scalebar = 2 μm. (e) Normalized ChIP-seq profile of NAT10 enrichment with active enhancer marks (H3K27ac and H4K20ac) in 4T1 cells. (f) IGV plot showing the ChIP enrichment of NAT10 with members of the nuclear pore-associated mechanosensitive protein complex at the *Myc* super-enhancer region. BRD4_short_V5 = V5-tagged short isoform of BRD4, RRP1B_HA = HA-tagged RRP1B, RRP1B_endo = endogenous RRP1B.

### NAT10 depletion affects transcription of chemokine and cytokine genes

Since NAT10 depletion promotes p300 mislocalization and disrupts enhancer organization, we next asked whether NAT10 depletion affects gene transcription. RNA-seq analysis revealed that NAT10 depletion had a robust effect on gene transcription. 783 genes were downregulated (Log2 fold change), and 1410 genes were upregulated (Log2 fold change) in *Nat10* KO 4T1 cells (Fig. 7a). Many of these top downregulated genes were chemokines (*Cxcl1*, *Cxcl2*, *Cxcl3*, *Cxcl5*, *Cxcl10*, *Cxcl16*, and *Ccl2*) and cytokine molecules (*Il1a*, *Il23a*, and *Tgfb2*). Some of these genes (*Cxcl1*, *Cxcl3*, *Ccl2*, and *Il1a*) were previously identified as NUP210-dependent mechanosensitive genes^8^ which supports a role of NAT10 in cellular mechanosensation. Overlaying the *Nat10* KO RNA-seq data with the H3K27ac ChIP-seq data revealed that 248 genes experienced a decrease of both the H3K27ac signal and mRNA expression (Fig. 7b). In contrast, 217 genes had a higher H3K27ac signal and increased expression. This result suggested that transcriptional alteration in a large set of genes in *Nat10* KO cells is due to the direct effect on enhancer organization.

**Fig. 7.**
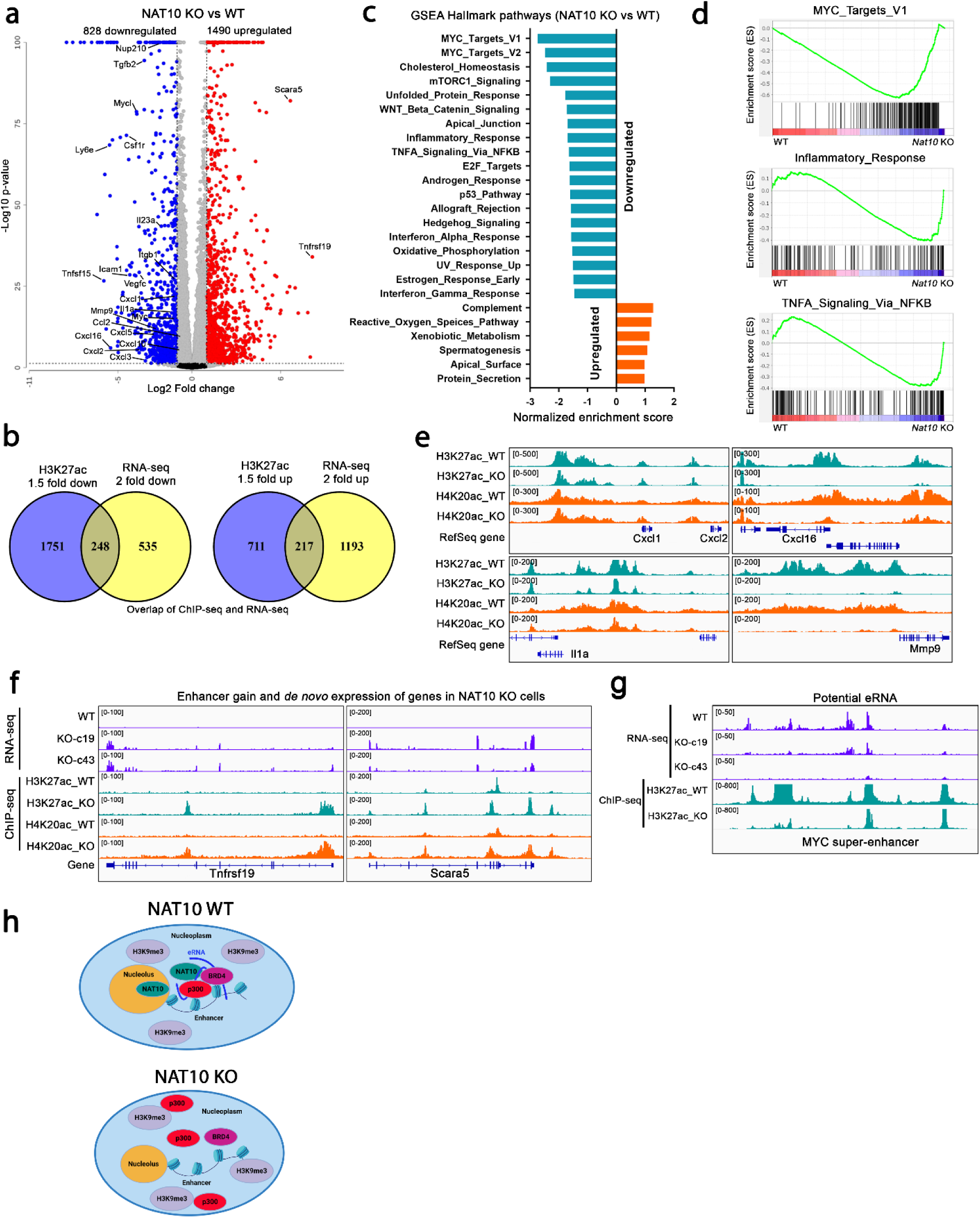
NAT10 depletion suppresses the transcription of chemokines and cytokine genes. (a) Volcano plot showing the distribution of up-and downregulated (Log2 fold change) genes in *Nat10* KO 4T1 cells. (b) Venn diagram showing the overlap of genes that have decreased H3K27ac enhancer marks and decreased transcription in *Nat10* KO 4T1 cells. (c) GSEA hallmark pathways down- and upregulated in *Nat10* KO 4T1 cells. (d) Example of key GSEA hallmark pathways downregulated in *Nat10* KO 4T1 cells. (e) IGV plot showing the decreased enrichment of enhancer marks (H3K27ac and H4K20ac) in the promoter of key immune-related, inflammatory response genes in *Nat10* KO 4T1 cells. (f)) IGV plot showing the genes that have gained enhancer marks and achieved *de novo* expression in *Nat10* KO 4T1 cells. (g) IGV plot showing the decreased level of potential enhancer RNAs (eRNA) within the *Myc* super-enhancer region in *Nat10* KO 4T1 cells. (g) Proposed model for NAT10 regulation of p300 localization within the nucleus.

GSEA gene ontology database analysis of *Nat10* KO cells revealed ‘Ribosomal Biogenesis’ as the top downregulated pathway (Extended Data Fig. 6a, b) suggesting a role of NAT10 in ribosomal biogenesis. In contrast, ‘Mammary Gland Epithelial Development’ and ‘Negative Regulation of Inflammatory Response’ were top upregulated pathways in *Nat10* KO 4T1 cells. The top 25 downregulated ribosome biogenesis genes in *Nat10* KO cells were mainly RNA polymerase II-dependent genes (Extended Data Fig. 6c). A moderate decrease in expression and enhancer (H3K27ac and H4K20ac) marks was found in the promoter regions of these genes (Extended Data Fig. 6c, d). As an aberrant increase in nucleolar size is reflective of increased ribosomal biogenesis^45^, we next examined nucleolar size in *Nat10* KO 4T1 cells. Although total nuclear size was decreased in *Nat10* KO cells, nucleolar area was not significantly changed (Extended Data Fig. 6e, f). Interestingly, nucleolar area was significantly decreased in p300 inhibitor (A-485)-treated cells (Extended Data Fig. 6g, h) indicating that p300 inhibition may also affect ribosome biogenesis. To assess whether the effect of NAT10 on p300 distribution is mediated by rRNA-dependent mechanism, we inhibited RNA polymerase I-dependent rRNA transcription by treating 4T1 cells with either BMH-21 (an RNA polymerase I-specific inhibitor) or low-dose actinomycin D (25 ng/ml, polymerase I inhibitory dose), and analyzed the distribution p300. Unlike the *Nat10* KO condition, treatment with polymerase I-specific inhibitors significantly decreased the p300 condensate intensity (Extended Data Fig. 6i, j), although residual p300 condensates significantly localized in the H3K9me3 heterochromatin zone. These results suggest that the observed effect of NAT10 depletion on ribosome biogenesis genes might be due to p300-dependent alteration of RNA polymerase II-mediated transcription.

To identify additional pathways that might contribute to the metastasis phenotype in *Nat10* KO cells, we performed GSEA hallmark pathway analysis. ‘MYC targets’ was one of the top hallmark pathways downregulated in *Nat10* KO cells (Fig. 7c, d), which is consistent with the decreased enhancer marks observed at the *Myc* super-enhancer region. In addition, some immune-related pathways such as ‘inflammatory response’, ‘TNFA signaling via NFκB’ were also downregulated in *Nat10* KO cells. Consistent with the downregulation, these chemokine/cytokines (*Cxcl1*, *Cxcl2*, *Cxcl16*, *Il1a*) as well as matrix metalloproteinase 9 (*Mmp9*) genes had decreased H3K27ac and H4K20ac enhancer marks in *Nat10* KO cells (Fig. 7e). Furthermore, genes such as *Tnfrs19* and *Scara5*, which gained enhancer marks, are typically silent in WT 4T1 cells but initiated *de novo* expression in *Nat10* KO cells (Fig. 7f). This may be driven by the mislocalization of p300 in developmentally silent heterochromatin regions in *Nat10* KO cells. As enhancers were affected by NAT10 depletion, we checked whether NAT10 depletion also affected enhancer RNAs (eRNA) expression. eRNAs are short RNA molecules transcribed from the enhancer regions which facilitate gene expression via facilitating enhancer condensates formation, recruiting chromatin-modifiers, and stimulating histone acetylation^46^. In the *Myc* super-enhancer region, potential eRNAs were expressed in WT 4T1 cells but their expression was decreased in *Nat10* KO cells (Fig. 7g) supporting a role of NAT10 in eRNA transcription. Therefore, we propose a model where NAT10 facilitates proper localization of p300 in enhancer condensates either through direct association or possible interaction with noncoding enhancer RNAs (eRNAs) in WT tumor cells (Fig 7h). Depletion of NAT10 leads to p300 mislocalization into heterochromatin regions, disruption of enhancer organization, and suppression of gene transcription.

### NAT10 depletion reshapes the innate immune tumor microenvironment and impairs neutrophil recruitment

Among the inflammatory and immune response genes downregulated in *Nat10* KO cells were the genes encoding for immune cell chemoattracting proteins (e.g., *Ccl2*, *Cxcl1*, *Cxcl2*, *Cxcl3*, *Cxcl5*, *Cxcl10*, and *Cxcl16*) (Fig. 2i, 7a). This prompted us to investigate the potential extracellular role of NAT10 in the regulation of the primary tumor microenvironment (TME). We found that immunocompromised mice lacking T and B lymphocytes injected with 4T1 *Nat10* knockdown cells continued to suppress tumor and lung metastasis (Extended Data Fig. 7a, b), suggesting a mechanism independent of the adaptive immune response. Therefore, we performed multiparametric flow cytometry to characterize the immune infiltration of 4T1 *Nat10* KO tumors implanted into BALB/c mice and the potential impact on the innate immune compartment. NAT10 depletion did not alter primary tumor growth (Extended Data Fig. 7c) or the total number of infiltrating leukocytes (CD45^+^) in the TME (Fig. 8a). However, we found substantial changes in the distribution of immune populations as shown by the unbiased t-Distributed Stochastic Neighbor Embedding (tSNE) projection of this single-cell data (Fig. 8b). Using unsupervised clustering followed by supervised cell annotation based on surface protein expression (Supplementary Table 2) we identified T and B adaptive lymphocytes and the following innate immune cell populations: neutrophils (Neu), eosinophils (Eos), mononuclear phagocytes (MPs), and innate lymphoid cells (ILCs) (Fig. 8b). *Nat10* KO tumors displayed significantly reduced neutrophil content, the most abundant immune cell type in 4T1 tumors, (Fig. 8b, c, d and Extended Data Fig. 7d, e). This is consistent with the transcriptomic data showing that the neutrophil-attracting chemokines (*Cxcl1*, *Cxcl2*, and *Cxcl5*)^47^ were reduced by NAT10 loss (Fig. 2i, 7a). We next focused our analysis on the second most abundant subset of innate immune cells in 4T1 tumors, MPs, which encompass monocytes (Mono), macrophages (Macs), and dendritic cells (DCs). The MP subset of cells was extracted, reclustered to define sub-populations, and visualized using Uniform Manifold Approximation and Projection (UMAP) that allows for increased spatial resolution to identify similarities between closely related cell clusters based on distance. We identified four clusters of Mono, five Macs, and four DCs whose proportions were dysregulated in *Nat10* KO resulting in an overall reduction of Mono and increase in Macs and DCs (Fig. 8e, f, Extended Data Fig. 7f, g). However, only total numbers of Mono were reduced by *Nat10* depletion (Fig. 8g, Extended Data Fig. 7h, i). This reduction was driven mainly by the Mono1 subset, which resemble classical Ly6C+ monocytes commonly found in circulation and is in line with the observed downregulation of their primary recruiting chemokine CCL2^48^ in *Nat10* KO cells (Fig. 2i, 7a and Extended Data Fig. 7h, i). Furthermore, analysis of Mac subsets divided into pro- or anti-tumorigenic phenotypes based on CD206 (pro-tumor associated) and MHCII (anti-tumor associated) expression showed an increase in the proportion of anti-tumor clusters Mac1 and Mac2 resulting in an overall reduction in the pro/anti-tumor Mac ratio in *Nat10* KO tumors (Fig. 8f, h, i, Extended Data Fig. 7g, i). These results indicate that the loss of NAT10 suppresses metastasis in part by shaping the innate immune compartment of the primary tumor microenvironment. Supporting our observations, several studies have shown a role for neutrophils^49–52^ and monocytes/macrophages^53, 54^ in driving 4T1 metastasis.

**Fig. 8.**
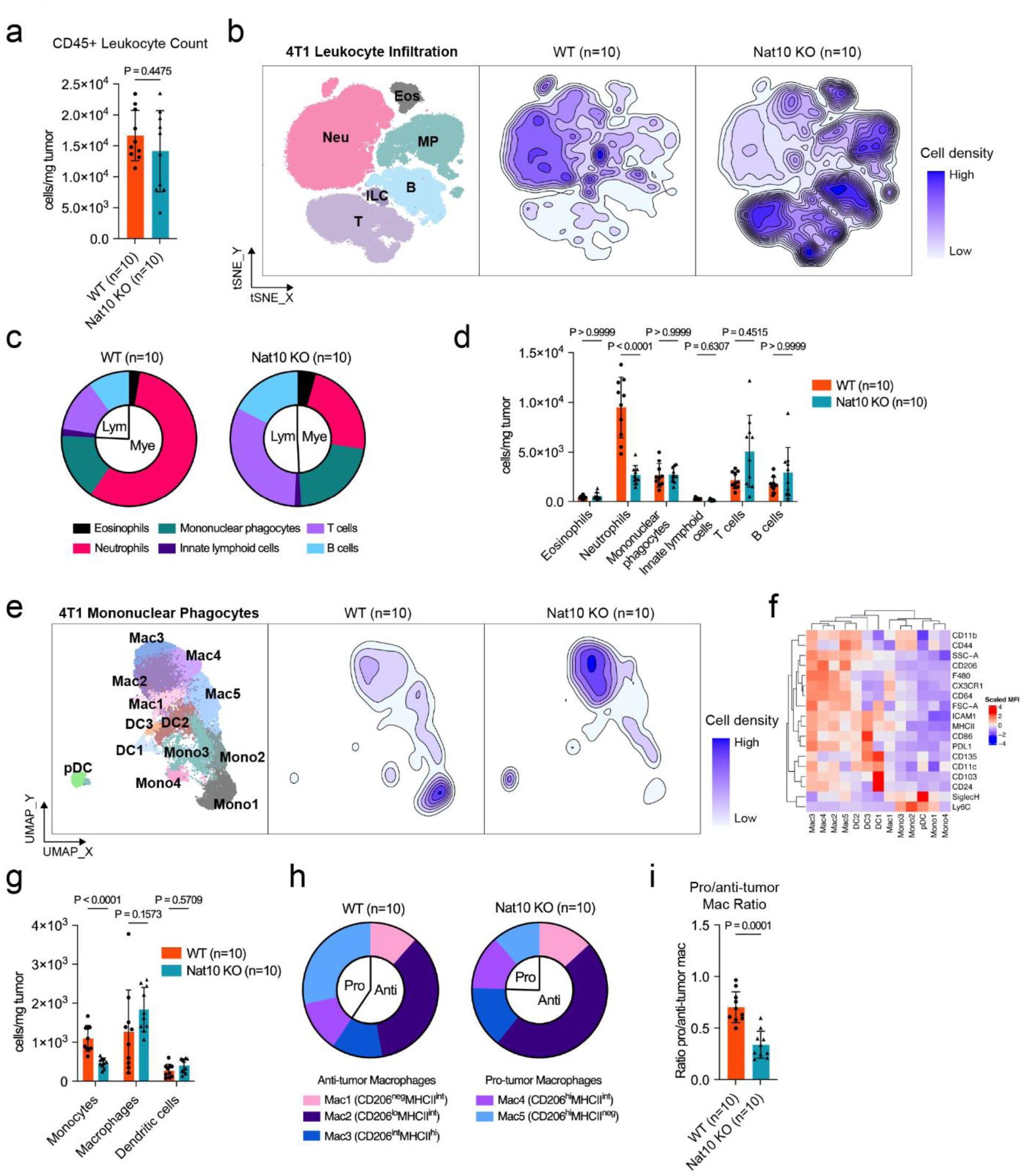
NAT10 depletion reduces neutrophils, monocytes, and pro-tumor macrophages in the tumor microenvironment. Tumor infiltrating leukocytes analyzed by high-parametric flow cytometry from 4T1 WT and *Nat10* KO-c43 primary tumors. (a) Absolute number of leukocytes (live CD45^+^) per mg of tumor. (b) t-SNE projection of tumor-infiltrating leukocytes; clustering of main immune cell populations (left): neutrophils (Neu), eosinophils (Eos), mononuclear phagocytes (MP), B cells (B), innate lymphoid cells (ILC) and T cells (T); contour density plots showing cell distribution in WT (center) and Nat10 KO-c43 (right) tumors. (c) Pie chart of main immune cell populations. (d) Absolute number of each main immune cell population per mg of tumor. (e) UMAP projection of MP subsets; clustering of MP subsets (left): monocytes (Mono1, Mono2, Mono3, Mono4), macrophages (Mac1, Mac2, Mac3, Mac4, and Mac5), and dendritic cells [DC1, DC2, DC3, and plasmacytoid dendritic cells (pDC)]; contour density plots showing cell distribution in WT (center) and Nat10 KO-c43 (right) tumors. (f) Heatmap representing the level of expression of indicated markers (scaled by marker) for each MP subset with hierarchical clustering indicated on top and markers on left. (g) Absolute number of each major MP population per mg of tumor. (h) Pie chart of macrophage subsets. (i) Pro/antitumor macrophage ratio. (a, i) Mann-Whitney U test, mean ± s.d. (d, g) Multiple Mann-Whitney test with Bonferroni-Dunn’s multiple comparisons test, mean ± s.d. n = 10 animals per group.

Taken together, our findings suggest that in response to the stiffness of the tumor microenvironment, *Nat10* expression in tumor cells facilitates transcription of metastasis-associated genes including chemokines and cytokines in a p300-dependent manner. These chemoattractant molecules then recruit neutrophil and monocytes, which promote metastatic dissemination of tumor cells from the primary tumor. Depletion of NAT10 leads to p300 mislocalization and disruption of enhancer organization. This ultimately leads to suppression of genes necessary for metastatic progression of mammary tumor cells along with genes for myeloid cell-attracting chemokines resulting in a less metastasis-prone primary tumor microenvironment.

## Discussion

In this study, we present evidence that Nat10 is a metastasis-associated factor for breast cancer. NAT10 was found to interact with the NUP210-bound mechanically sensitive protein complex at the nuclear pore that we recently described. We now demonstrate that NAT10 depletion phenocopies many of the cellular and *in vivo* phenotypes of NUP210 depletion, suggesting that NAT10 is an important component of NUP210-associated response to extracellular stiffness and cellular motility. Importantly, multiple independent screens identified components of this complex, namely SIPA1, BRD4, RRP1B, and NUP210. The repeated identification of these common pathway components by independent methods and their common phenotypic effects, such as cellular adhesion, suggests that this mechanically responsive pathway plays an important role in tumor progression in breast cancer. Since breast cancer cells begin to disseminate early in tumor evolution as tumors start to become hypoxic and fibrotic^55^, this pathway is most likely associated with the dissemination phase of the metastatic cascade. Further investigations into this expanding pathway may therefore provide important insights into the etiology of metastasis.

NAT10 can function as an RNA cytidine acetyltransferase, and is suggested to acetylate various proteins, including histones and p53^15–18^. NAT10 has also been implicated in promoting metastasis in multiple cancer types including breast cancer^25, 27, 56–58^. However, most of these studies have focused on effects on individual mRNAs. Although the key RNA or protein target of NAT10 in metastatic breast cancer is currently unclear, here we demonstrate that a 50% reduction of NAT10 acetyltransferase activity through the expression of an endogenous acetylase-inactivated allele in the MMTV-PyMT model significantly reduces metastatic capacity. Similar results were observed in both shRNA depletion and sgRNA knockout studies, supporting an important role of NAT10 in metastatic progression. However, rather focusing on single gene downstream of NAT10, our findings implicate NAT10 in the regulation of chromatin architecture and transcriptional control. Similar to NUP210-depleted cells, NAT10 depletion resulted in significant nuclear rearrangement and sequestration of histone 3.1/3.2 from the nuclear periphery to heterochromatin foci. In addition, consistent with its proposed function as a histone acetyltransferase, NAT10-depletion resulted in a marked decrease in histone acetylation. *In vitro* acetylation assays, however, did not support a role for NAT10 in histone or p300 acetylation. While we cannot dismiss the possibility that the *in vitro* assay conditions did not properly support NAT10 histone acetylation activity, we lean toward attributing the decrease in histone acetylation to indirect effects rather than direct acetylation effects. Further research will be required to clarify the metastasis-associated acetylation targets of NAT10.

Unexpectedly, the *in vitro* acetylation assays led to the discovery that NAT10 interacts with p300 and can modulate p300 function and subnuclear localization. Our results suggest that NAT10 is an obligatory co-factor of p300 transcriptional regulatory network and is likely required for proper p300 localization on chromatin. However, how NAT10 regulates p300 localization remains unclear. One possibility is that NAT10 serves as a scaffolding protein to directly facilitate p300 localization on enhancers. Alternatively, p300 localization on enhancers may be indirectly regulated by NAT10-mediated acetylation and stabilization of non-coding RNAs. Evidence suggests that p300 can be recruited to many enhancer sites by the long non-coding RNA (lncRNA)^59^, and NAT10 has been implicated in lncRNA acetylation that facilitates metastasis^56^. Therefore, it is possible that NAT10 catalyzes the acetylation of an unknown LncRNA, which then facilitates the proper enhancer localization of p300. Moreover, CBP, a p300 paralog, has been shown to bind to relatively short, non-coding, enhancer RNAs (eRNAs)^60^. Binding CBP to these eRNAs can stimulate its HAT activity. Here, we showed that potential eRNAs transcribed from the *Myc* super-enhancer region were downregulated in *Nat10* KO cells. eRNAs have been shown to facilitate transcriptional condensate formation^46, 61^. NAT10 may influence the transcription or catalyze ac4C-modification of these eRNAs, potentially impacting p300 localization at transcriptional condensates. Future studies should determine whether ac4C modification of eRNA alters transcriptional condensate formation.

As anticipated due to the disruption of p300 histone acetyltransferase activity, the depletion of NAT10 led to significant changes in cellular transcription. Interestingly, NAT10 loss decreases the expression of immune-related chemokines and cytokines. However, metastasis suppression persisted when NAT10-depleted cells were orthotopically transplanted into nude mice, arguing against a role for the adaptive immune response. Conversely, we found that NAT10 KO primary tumors have a remodeled innate tumor microenvironment marked by a reduction in neutrophils and monocytes, likely due to reduced production of key myeloid-attracting chemokines such as CCL2, CXCL1, CXCL2, CXCL3, CXCL5, and a reduced pro-tumorigenic macrophage phenotype. Through different approaches such as antibody depletions^49^, inhibiting recruitment via CXCL1/CXCL2 supression^50^, and blocking granulocyte-colony stimulating factor (G-CSF)^51^, several studies have established the role of neutrophils in promoting breast cancer lung metastasis^52, 62^. Other studies have additionally implicated monocytes and/or macrophages in breast cancer metastasis^53, 54, 63^. Previously we found that overexpression of *Ccl2* in *Nup210* KO cells incompletely restored lung metastatic burden to control levels^8^. Considering that NAT10 loss phenocopies several NUP210-dependent pathways, it is likely that CCL2 targeting only restored monocyte recruitment whereas neutrophil recruitment was still impaired. Simultaneous targeting of CCL2 along with neutrophil-targeting chemoattractant CXCL1 and CXCL2 may be required to sufficiently restore the metastasis phenotype. Alternatively, there could be downstream targets of NAT10/NUP210 that are independently necessary for tumor metastatic potential in conjunction with extracellular communications with myeloid cells. Further studies are needed to fully dissect the interplay between tumor NAT10 expression and intratumoral/systemic myeloid cells governing metastatic progression.

The role of NAT10 in super-enhancer function also raises a potential for clinical applications. Bromodomain and extra-terminal domain (BET) protein inhibitors such as JQ1 and I-BET151 were originally designed to suppress *Myc* oncogene expression by disrupting the binding of BRD4 to the *Myc* super-enhancer. The reduction of NAT10 function also suppresses MYC expression and the transcription of downstream targets. Therefore, designing a specific inhibitor for NAT10 may provide an orthogonal strategy to attenuate MYC activity in *Myc*-amplified cancers. Remodelin, a potential NAT10 inhibitor, normalizes nuclear morphology in cells from patients with laminopathies^40^. However, recent studies have suggested that it is not a specific NAT10 inhibitor since it does not inhibit the ac4C-modification of RNA molecules^64^. Development of additional, more specific inhibitors of NAT10 might provide alternative methods or combinatorial opportunities to improve patient outcomes through either adjuvant or anti-metastatic therapies.

In conclusion, NAT10 is a mechanically sensitive regulator of metastasis in breast cancer functioning through its association with the previously described mechanically sensitive microenvironmental sensor mechanism that involves the nuclear pore protein, NUP210. NAT10’s impact on metastasis occurs through a novel interaction with the histone acetyltransferase p300, establishing pro-metastatic chromatin structure and transcriptional programs. The loss of NAT10 activity leads to the reorganization of chromatin architecture and mislocalization of the p300 into heterochromatin foci, resulting in significant changes in transcription and cellular adhesion. The metastasis-promoting function of NAT10 is associated with its acetyltransferase activity, though the precise target in the context of metastasis remains known. Furthermore, NAT10 loss led to reduced production of myeloid cell recruiting chemokines, which reduced metastasis-promoting neutrophils, monocytes, and macrophages in the primary tumor. Further research will be required to elucidate the metastasis-associated acetylation target and to extensively characterize the role of NAT10 in regulating tumor-immune crosstalk and metastatic progression.

## Acknowledgements

Authors thank Brandi Carofino for critical review of the manuscript, Center for Cancer Research (CCR) Laboratory of Genome Integrity Flow Cytometry Core Facility Head Ferenc Livak, CCR Confocal Microscopy Core Facility Head Michael J. Kruhlak for microscopy analysis, CCR Genomics Core facility member Liz Conner, CCR sequencing facility (Frederick, MD) for RNA-seq/ChIP-seq analysis. The study was supported by the Intramural Research Program, National Cancer Institute, National Institutes of Health.

## Author contributions

R.A., K.W.H., and N.H. conceived the project. R.A performed majority of the experiments, analyzed the data, and wrote the manuscript. N.H. performed *in vivo* mouse experiment, OPP assay and analyzed the data. T.Q. performed mouse breeding and analyzed the data. K.C.L. and A.L. performed the immunological experiment, analyzed the data, and wrote the manuscript. R.H. performed mass spectrometry analysis, analyzed the data, and wrote the manuscript. H.L. and M.P.L. performed bioinformatics analysis. A.D.T. performed microscopy image analysis data.

S.T.G performed ac4C analysis and wrote the manuscript. R.S.G. supervised the immunological analysis and wrote the manuscript. J.L.M. supervised the ac4C analysis and wrote the manuscript.

T.A. supervised the mass spectrometry analysis. K.W.H. wrote the manuscript and supervised the whole project.

---

**Extended Data Fig. 1.**
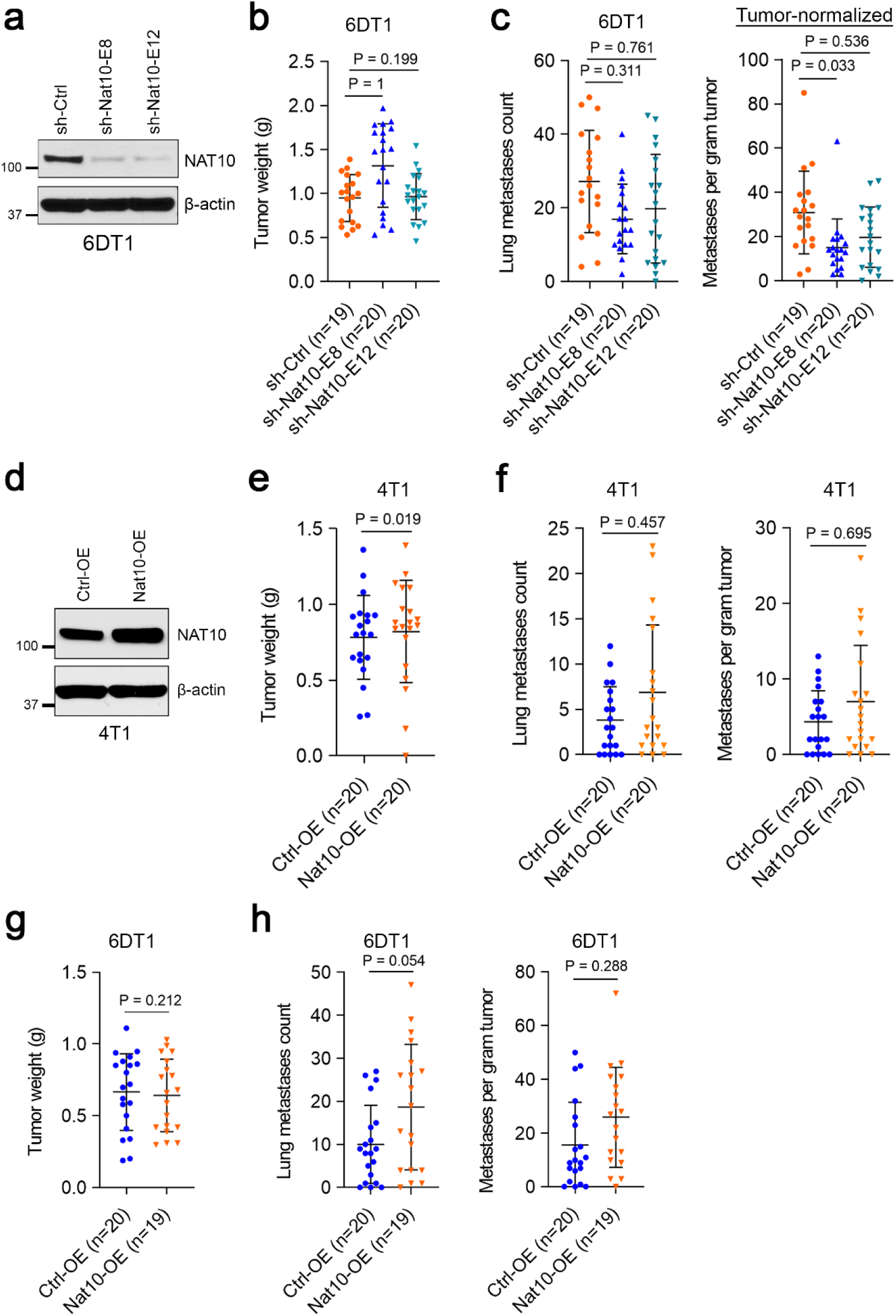
**Related to Fig.1**. (a) Western blot showing the level of NAT10 protein in shRNA knockdown 6DT1 cells. (b) Tumor weight of mice injected with *Nat10* knockdown 6DT1 cells. Kruskal-Wallis ANOVA with Dunn’s multiple comparison test, mean ± s.d. p-values from two independent experiments were combined with Fisher’s method. (c) Lung metastases count (left) and tumor-normalized metastases count (right) in *Nat10* shRNA knockdown 6DT1 cells. Kruskal-Wallis ANOVA with Dunn’s multiple comparison test, mean ± s.d. p-values from two independent experiments were combined with Fisher’s method. (d) Western blot showing the overexpression of NAT10 protein in 4T1 cells. (e) Tumor weight of mice injected with *Nat10*-overexpressing 4T1 cells. Mann-Whitney U test, mean ± s.d. p-values from two independent experiments were combined using Fisher’s method. (f) Lung metastases count (left) and tumor-normalized lung metastases count (right) in mice injected with *Nat10*-overexpressing 4T1 cells. Mann-Whitney U test, mean ± s.d. p-values from two independent experiments were combined using Fisher’s method. (g) Tumor weight of mice injected with *Nat10*-overexpressing 6DT1 cells. Mann-Whitney U test, mean ± s.d. p-values from two independent experiments were combined using Fisher’s method. (h) Lung metastases count (left) and tumor-normalized lung metastases count (right) in mice injected with *Nat10*-overexpressing 6DT1 cells. Mann-Whitney U test, mean ± s.d. p-values from two independent experiments were combined using Fisher’s method.

**Extended Data Fig. 2.**
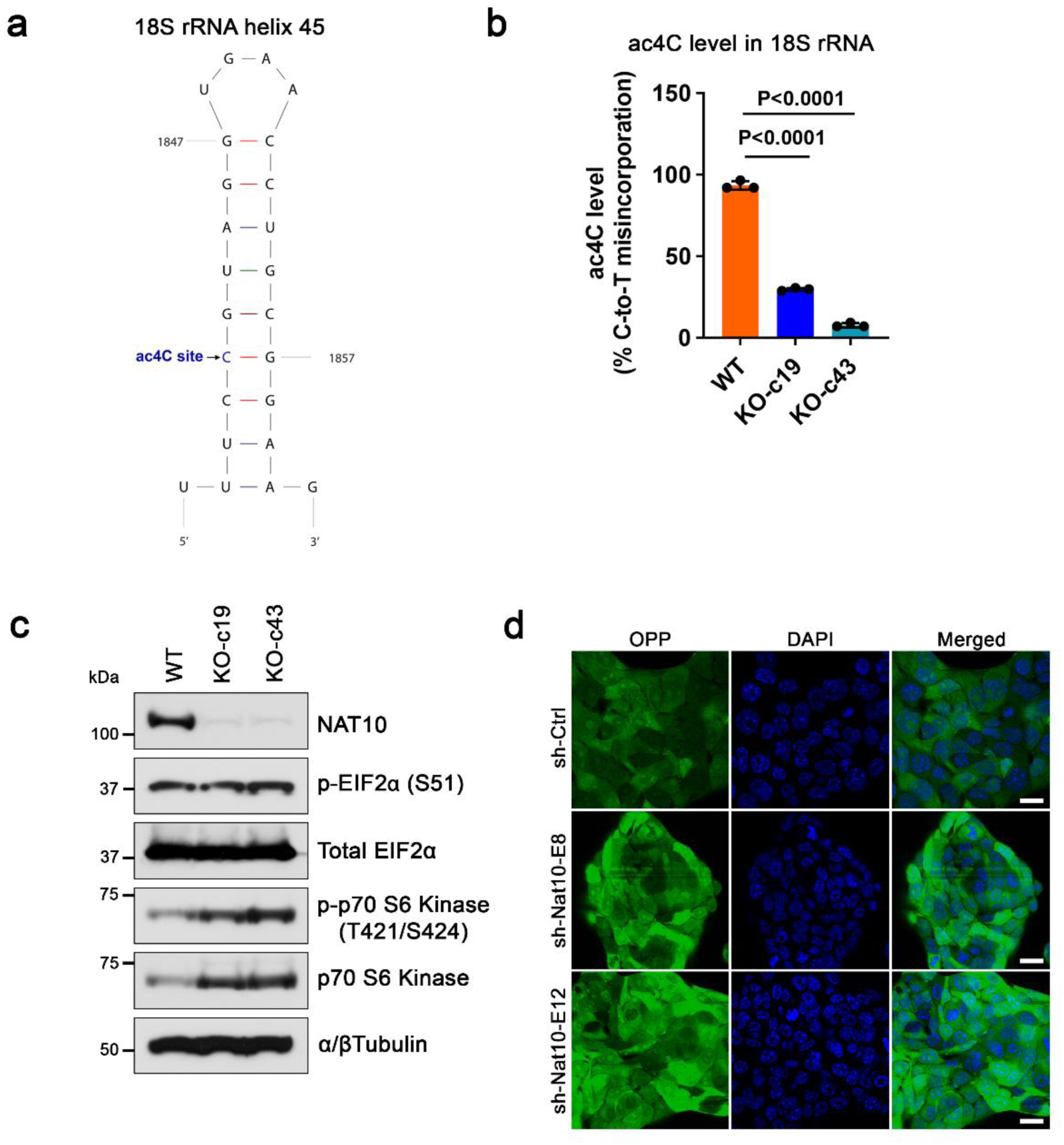
Effect of Nat10 depletion on ac4C modification and translational control pathway components. (a) ac4C modification site of 18S rRNA helix 45. (b) ac4C modification analysis of 18S rRNA helix 45 in *Nat10* KO 4T1 cells. ANOVA with Tukey’s multiple comparison test, mean ± s.d. (c) Western blot showing the level of translational control pathway components in *Nat10* KO 4T1 cells. (d) OPP assay showing the level of protein production in *Nat10* knockdown 4T1 cells. Scale bar = 10 μm.

**Extended Data Fig. 3.**
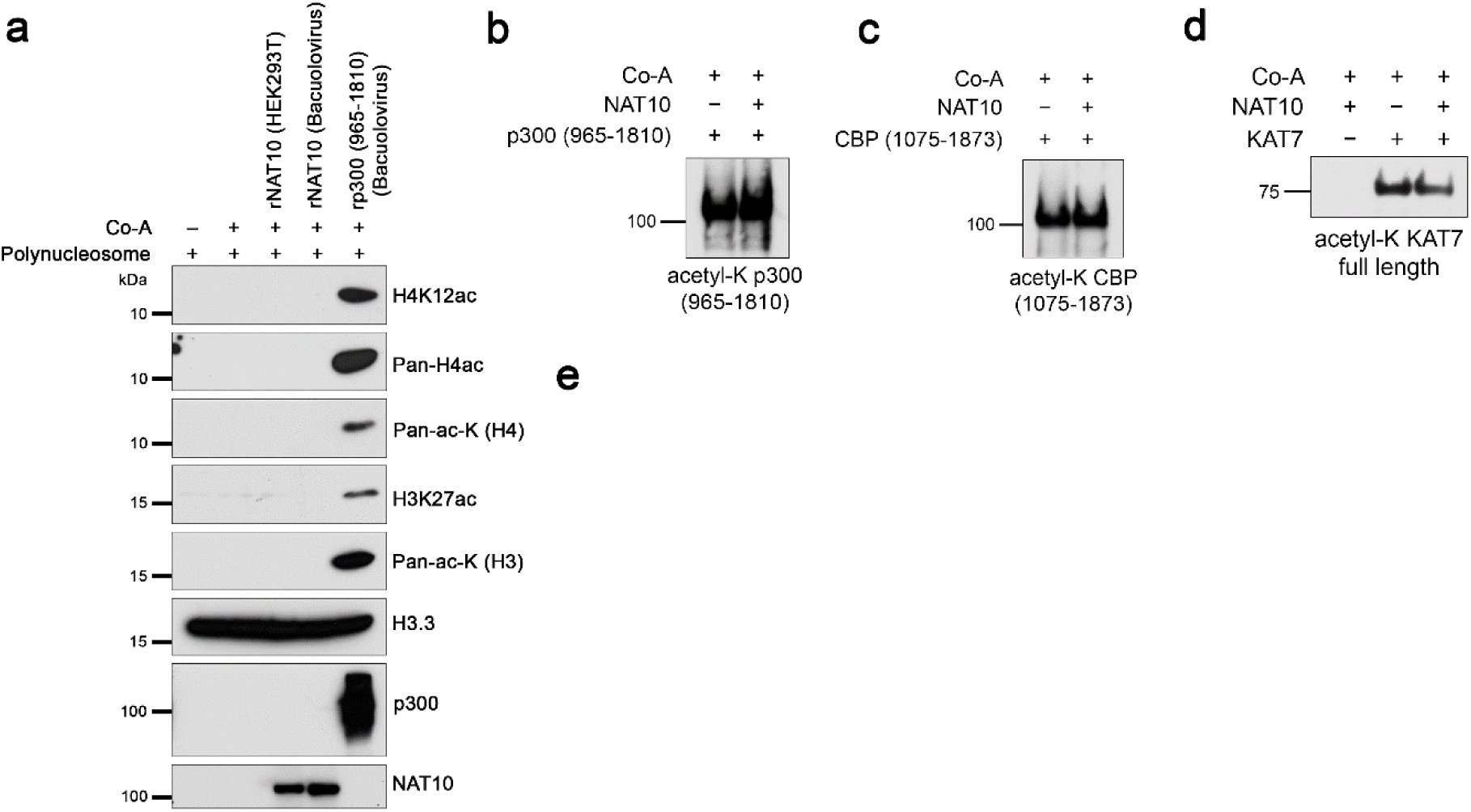
**Related to Fig. 3**. (a) Western blot of the *in vitro* HAT assay with polynucleosome, recombinant NAT10 and p300 catalytic subunits. (b) Western blot of the *in vitro* HAT assay with recombinant NAT10 and p300 catalytic subunit. (c) Western blot of the *in vitro* HAT assay with recombinant NAT10 and CBP catalytic subunit. (d) Western blot of the *in vitro* HAT assay with recombinant NAT10 and KAT7.

**Extended Data Fig. 4.**
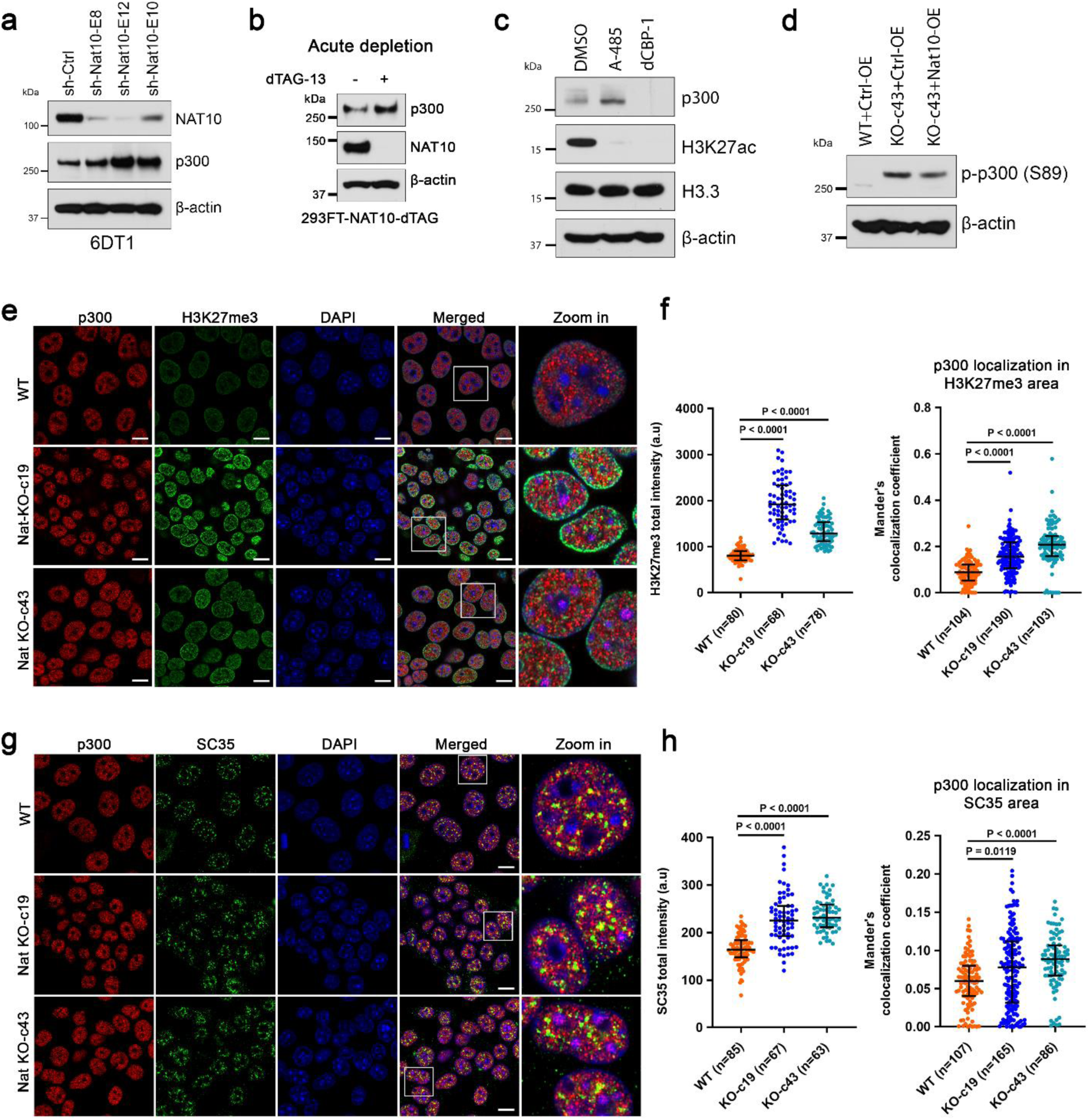
**Related to Fig.4**. (a) Western blot showing the level of p300 protein in *Nat10* shRNA knockdown 6DT1 cells. (b) Western blot showing the level of p300 protein after acute degradation of NAT10 in human 293FT cells. (c) Western blot showing the protein level of p300 and H3K27ac in A-485- and dCBP-1-treated 4T1 cells. (d) Western blot showing the protein level of phospho-p300 (S89) in *Nat10* KO and rescued 4T1 cells. (e) Immunofluorescence showing the distribution of p300 and H3K27me3 mark within the nuclei of *Nat10* KO 4T1 cells. Scalebar = 10 μm. (f) Quantification of H3K27me3 signal in *Nat10* KO 4T1 cells (left), Kruskal-Wallis ANOVA with Dunn’s multiple comparison test, median with interquartile range, Mander’s colocalization coefficient of p300 colocalization in the H3K27me3 region in *Nat10* KO 4T1 cells (right), Kruskal-Wallis ANOVA with Dunn’s multiple comparison test, median with interquartile range. (g) Immunofluorescence showing the distribution of p300 and nuclear speckle marker SC35 within the nuclei of *Nat10* KO 4T1 cells. (h) Quantification of SC35 signal in *Nat10* KO 4T1 cells (left), Kruskal-Wallis ANOVA with Dunn’s multiple comparison test, median with interquartile range, Mander’s colocalization coefficient of p300 colocalization in the SC35 region in *Nat10* KO 4T1 cells (right), Kruskal-Wallis ANOVA with Dunn’s multiple comparison test, median with interquartile range.

**Extended Data Fig. 5.**
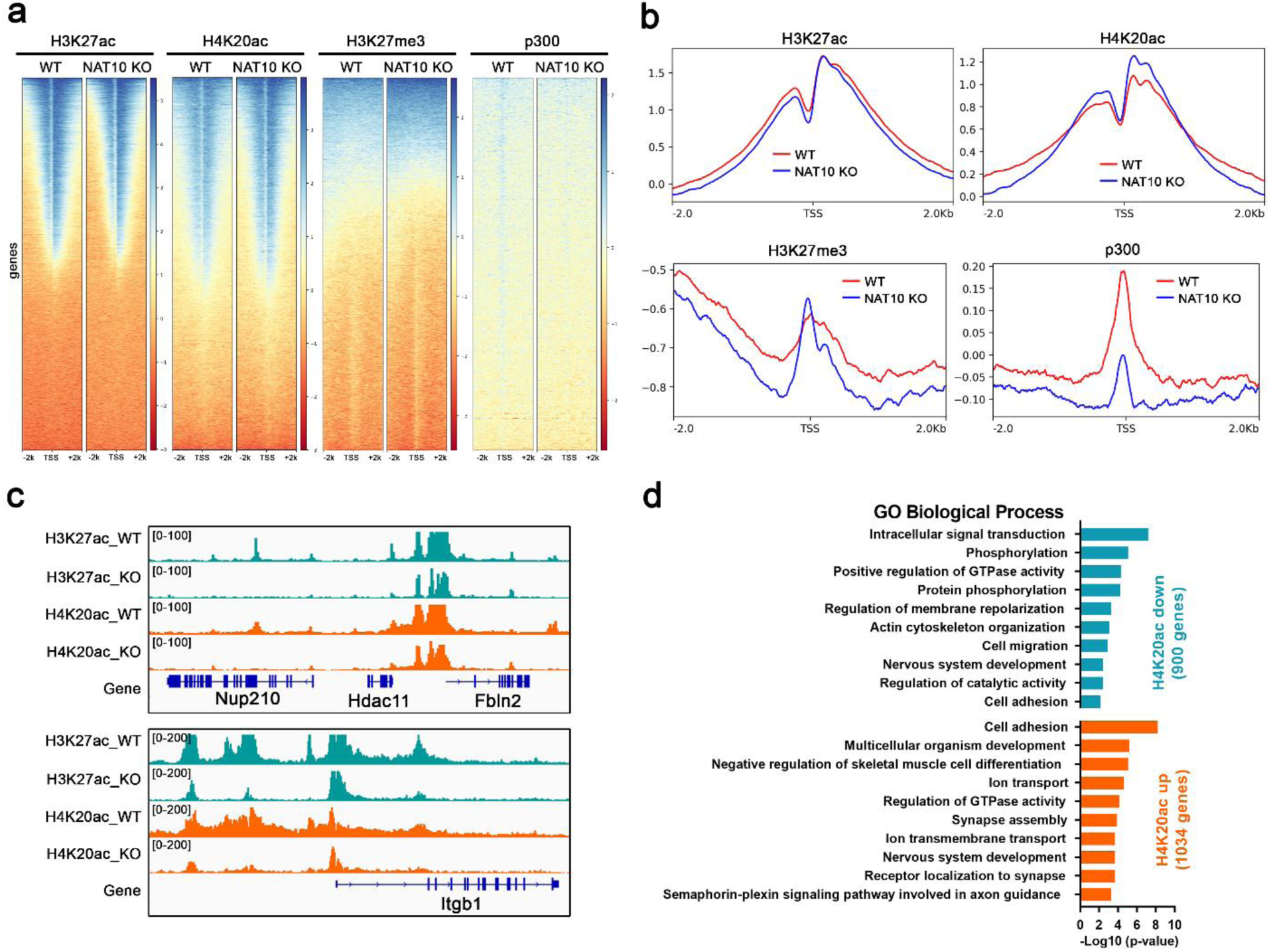
**Related to Fig.5**. (a) ChIP-seq heatmap showing the peak distribution of H3K27ac, H4K20ac, H3K27me3, and p300 within the transcription start sites (TSS) of *Nat10* KO 4T1 cells. (b) Normalized ChIP enrichment profile of H3K27ac, H4K20ac, H3K27me3, and p300 within the TSS of *Nat10* KO 4T1 cells. (c) IGV plot showing the decreased enrichment of H3K27ac and H4K20ac at cell adhesion genes (*Nup210* and *Itgb1*) loci in *Nat10* KO 4T1 cells. Gene Ontology pathway analysis for genes that either lost or gained H4K20ac marks in *Nat10* KO 4T1 cells.

**Extended Data Fig. 6.**
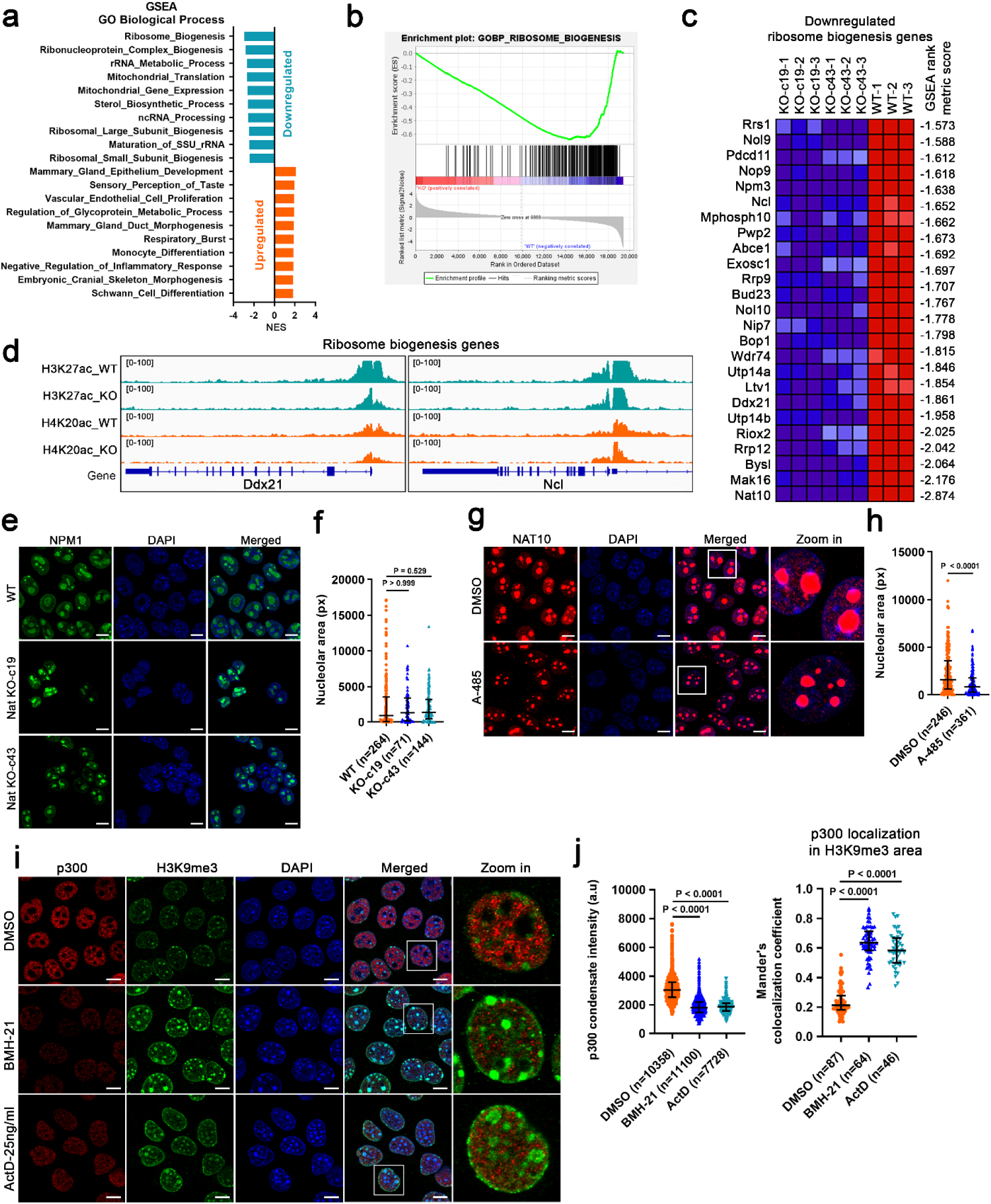
**Related to Fig.7**. (a) GSEA Gene Ontology pathway analysis of *Nat10* KO 4T1 cells. (b) Key Gene Ontology pathway enriched in *Nat10* KO 4T1 cells. (c) GSEA heatmap of the top 25 downregulated ribosome biogenesis genes in *Nat10* KO 4T1 cells. (d) Enrichment of H3K27ac and H4K20ac in the promoter region of key ribosome biogenesis genes. Immunofluorescence showing the morphology of NPM1-marked nucleoli in *Nat10* KO 4T1 cells, Scalebar = 10 μm. (f) Quantification of nucleolar area in *Nat10* KO 4T1 cells. Kruskal-Wallis ANOVA with Dunn’s multiple comparison test, median with interquartile range, px = pixel. (g) Immunofluorescence showing the morphology of NAT10-marked nucleoli in A-485-treated 4T1 cells, Scalebar = 10 μm. (h) Quantification of nucleolar area in A-485-treated nucleoli in 4T1 cells, Mann-Whitney U test, median with interquartile range, px = pixel (i) Immunofluorescence showing the distribution of p300 and H3K9me3 in BMH-21 and low dose actinomycin D (ActD)-treated 4T1 cells. Scale bar = 10 μm. (j) Quantification of p300 condensate intensity (left) and Mander’s colocalization coefficient showing p300 localization in H3K9me3 area (right) in BMH-21- and ActD-treated 4T1 cells. Kruskal-Wallis ANOVA with Dunn’s multiple comparison test, median with interquartile range.

**Extended Data Fig. 7.**
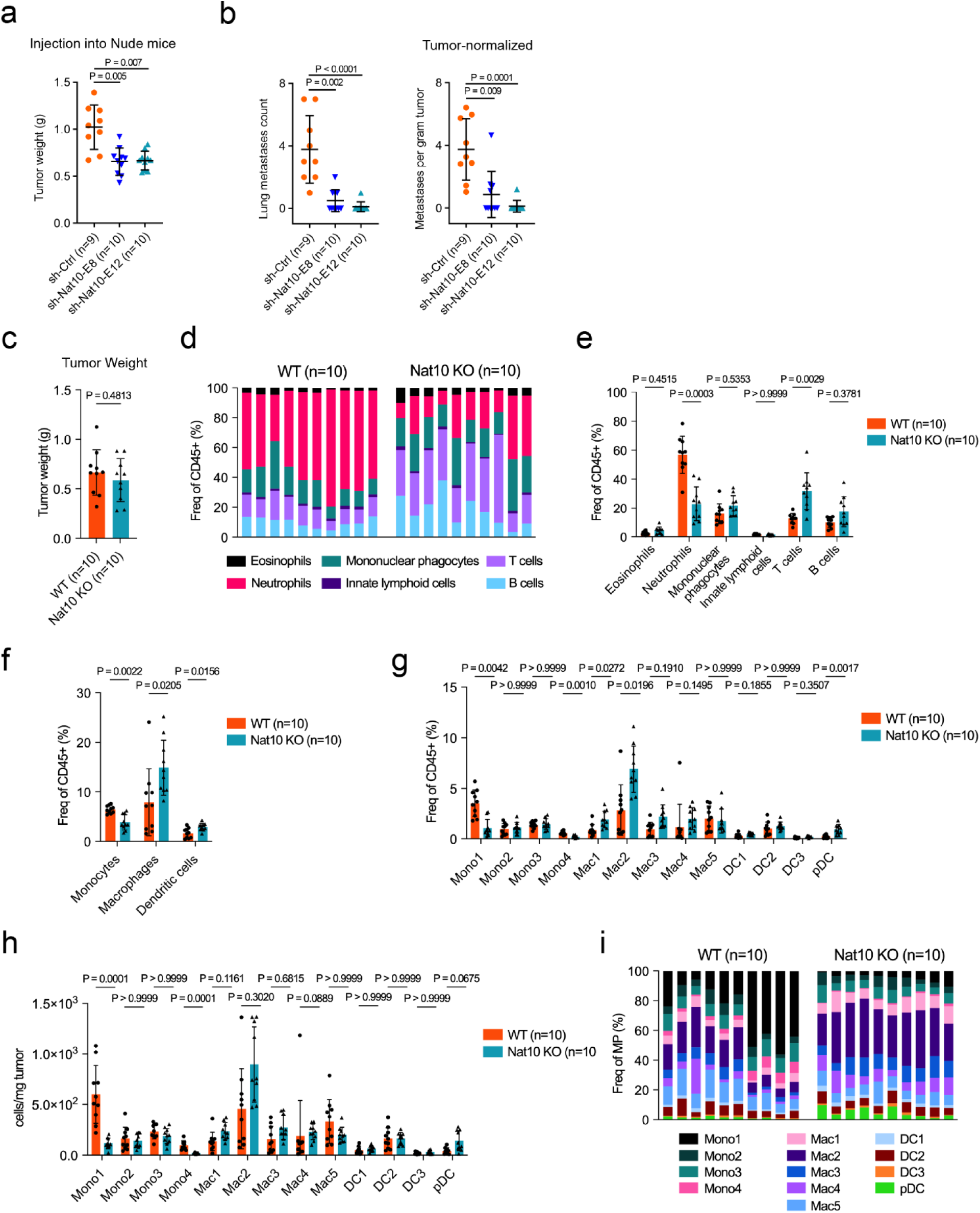
**Related to Fig.8**. (a) Primary tumor weight after injecting *Nat10* knockdown 4T1 cells in athymic nude mice. Kruskal Wallis ANOVA with Dunn’s multiple comparison test, mean ± s.d. (b) Lung metastases count (left), and tumor-normalized metastases count (right) after injection of *Nat10* knockdown 4T1 cells in athymic nude mice. Kruskal Wallis ANOVA with Dunn’s multiple comparison test, mean ± s.d. (c) 4T1 WT and *Nat10* KO-c43 primary tumor weight. (d-i) Tumor infiltrating leukocytes analyzed by high-parametric flow cytometry from 4T1 WT and Nat10 KO-c43 primary tumors. (d) Proportion of indicated leukocyte population in total leukocyte for each individual sample. (e) Proportion of indicated leukocyte population in total leukocyte. (f) Proportion of indicated mononuclear phagocyte (MP) population in total leukocyte. (g) Proportion of indicated MP subsets in total leukocyte. (h) Proportion of indicated MP subsets in total leukocyte for each individual sample. (i) Absolute number of each MP subsets per mg of tumor. (c) Mann-Whitney U test, mean ± s.d. (e, f, g, h) Multiple Mann-Whitney test with Bonferroni-Dunn’s multiple comparisons test, mean ± s.d. n = 10 animals per group.

**Supplementary Table 1:**
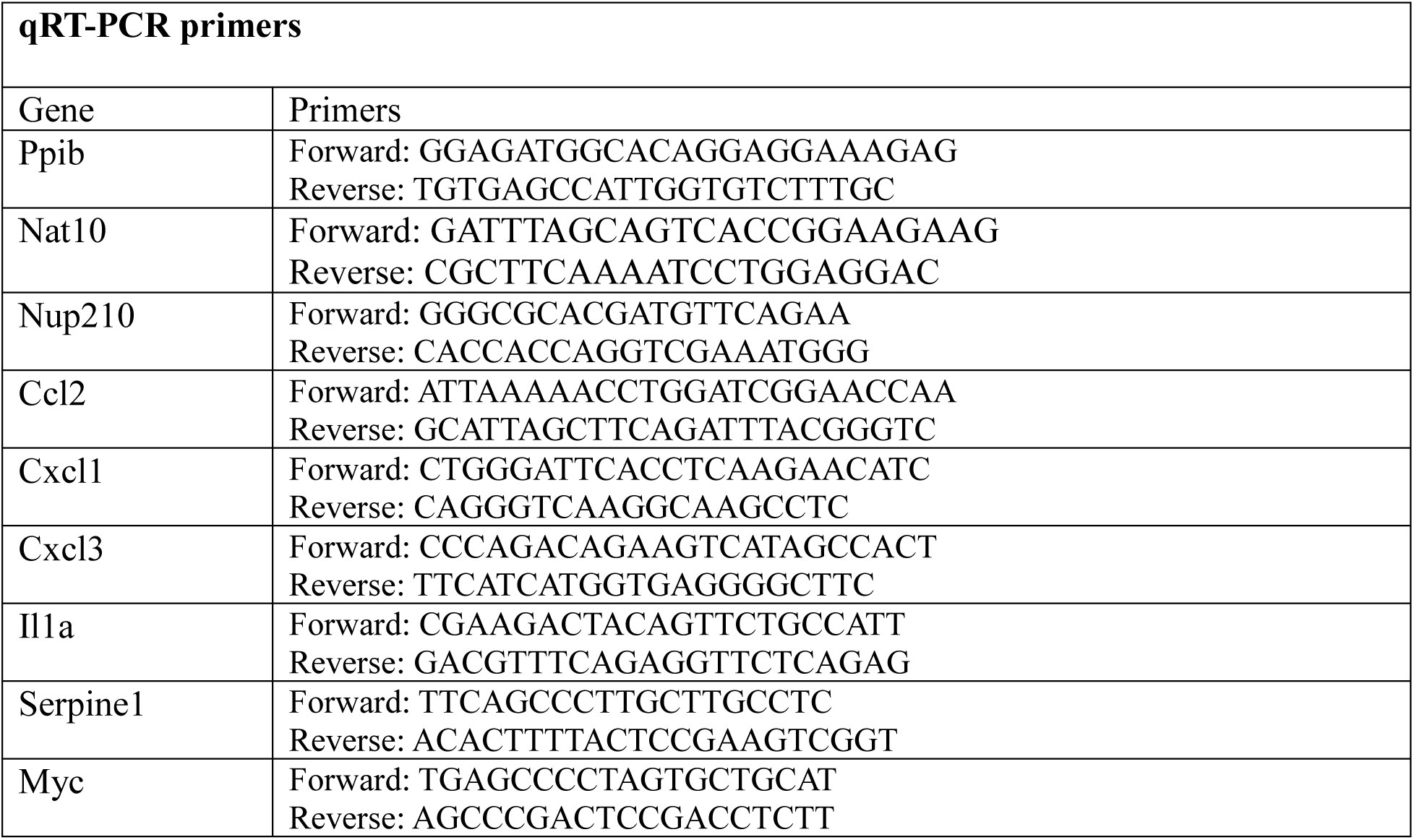
List of oligonucleotides used in this study:

**Supplementary Table 2:**
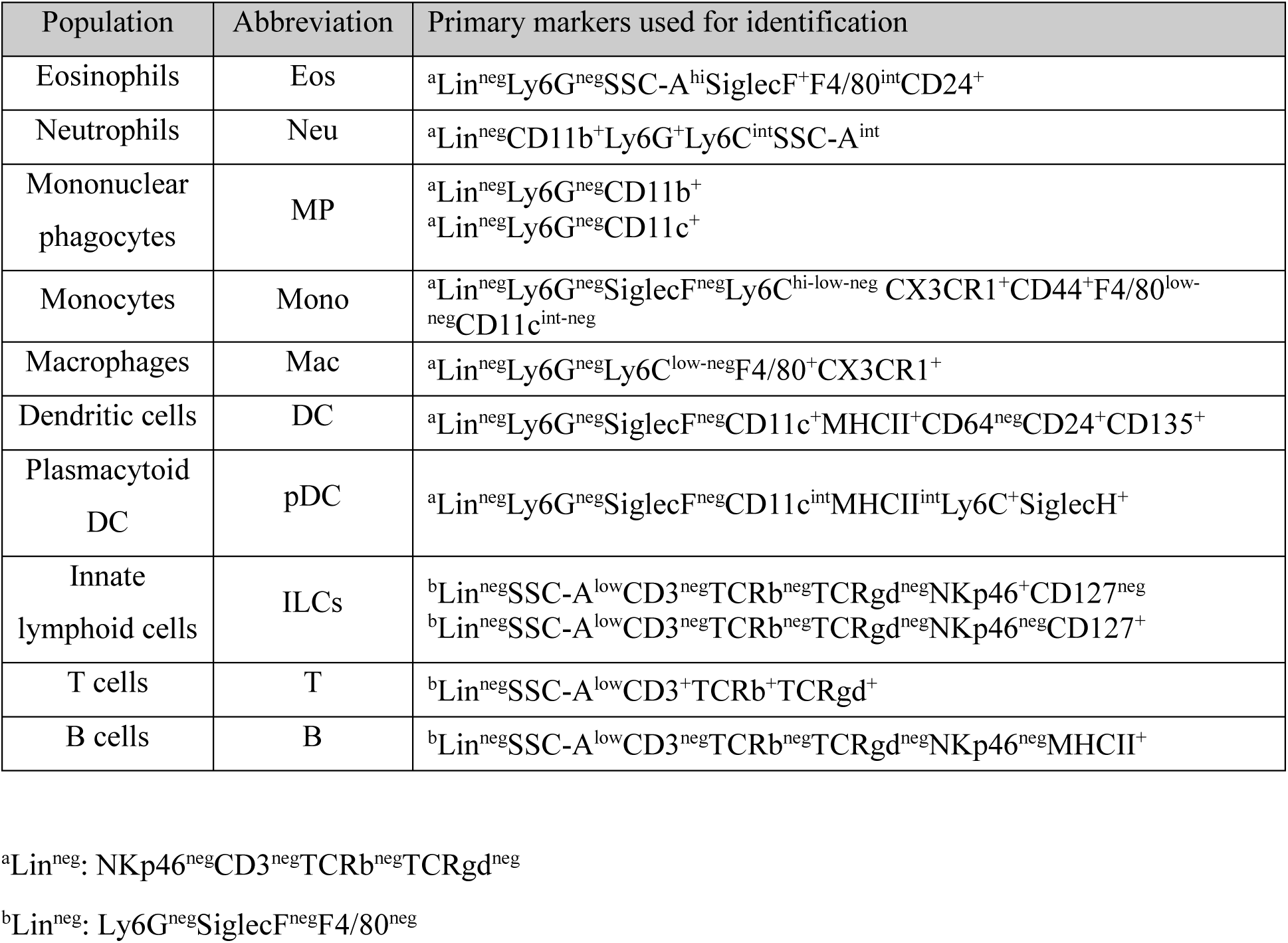

